# Ebola’s Hidden Target: Virus Transmission to and Accumulation within Skin

**DOI:** 10.1101/2025.04.04.647113

**Authors:** Paige T. Richards, Anthony Fleck, Radhika Patel, Maryam Fakhimi, Dana Bohan, Kathleen Geohegan-Barek, Allison E. Stolte, Caitlin O. Messingham, Samuel Connell, Tyler Crowe, Francoise Gourronc, Ricardo Carrion, Anthony Griffiths, David K Meyerholz, Al Klingelhutz, Robert A. Davey, Kelly N. Messingham, Wendy Maury

## Abstract

Ebola virus (EBOV), the causative agent of Ebola virus disease (EVD), remains one of WHO’s top ten threats to global health. Infectious EBOV virions can be found on the surface of skin late during systemic infection and passed from the deceased through skin-to-skin contact. Here, we assess viral load and antigen expression in the skin of EBOV-infected non-human primates (NHP) and mouse adapted-EBOV (ma-EBOV) - infected mice and use the low containment viral model, rVSV/EBOV GP, to mechanistically define skin infection in mice. Viral RNA peaked within the skin proximal to the site of injection in EBOV-infected NHPs on day 6. In contrast, mouse skin sites distal to the site of ma-EBOV injection achieved maximal viral loads by day 3. At late times of infection, viral antigen-positive cells co-localized with markers for endothelial, stromal, and immune cells in the dermis. Epidermal cells within and surrounding hair follicles also harbored viral antigen, suggesting a potential mechanism of virus trafficking to the epidermal surface. Despite robust viral infection, distal skin sites of ma-EBOV-infected mice had low expression of proinflammatory stimulated genes. A similar cellular tropism was observed in the skin of mice infected with rVSV/EBOV GP, with discrete focal areas of intense infection. When virus was applied to the surface of gently abraded skin to remove the stratum corneum, epidermal keratinocytes were robustly infected, followed by systemic viral dissemination. To define cell surface receptors critical for virus trafficking to and replication within the skin, mice lacking the phosphatidylserine receptors were infected intraperitoneally with rVSV/EBOV GP. At day 3 of infection, skin distal to the site of infection of TIM-1 knock out (KO) mice had significantly lower levels of infectious virus than the control mice, suggesting that TIM-1 is essential for efficient distribution of virus to the skin. Our findings reveal that EBOV targets specific skin cell populations at late times of viral infection and that the host receptor TIM-1 is required for optimal viral dissemination.

## INTRODUCTION

Sporadic Ebola virus (EBOV) outbreaks continue to occur in Central and Western Africa despite the availability of an FDA-approved vaccine (1). It is therefore not surprising why EBOV remains one of the WHO’s top ten global health threats (2). The largest epidemic, from 2013 to 2015 in West Africa, resulted in approximately 28,000 reported cases and over 11,000 deaths (1). EBOV infection can cause severe symptoms, including hemorrhagic fever, gastrointestinal distress, multiple organ failure, cytokine dysregulation, and death (3).

EBOV primarily spreads through direct contact with infected bodily fluids (3). Mucosal transmission is considered the main route of infection (3). However, during the West African outbreak, anecdotal evidence suggests infectious virus is present on the skin of patients with late-stage infections (4). This poses a significant risk of transmission via skin contact. Consistent with this, EBOV RNA and infectious virions are detectable on the skin in late-stage infection in animal models (5).

Our lab recently developed a transwell human skin explant model to investigate direct EBOV infection of skin (6). We demonstrated that EBOV infection via the basal dermal surface occurred in a time- and dose-dependent manner, trafficking through the tissue to the apical epidermal surface, where infectious virus or RNA was often detected. Dermal tropism included cells of myeloid, endothelial, epithelial, and stromal origin. We also assessed EBOV-specific dermal entry receptors. Along with other enveloped and members of the *Filoviridae* family, EBOV can enter cells via apoptotic mimicry (7). Using phosphatidylserine (PS) on its envelope, EBOV virions bind to one of several PS receptors—either a TIM or TAM receptor—inducing endocytosis. The viral glycoprotein (GP) is proteolytically processed within the endosomal compartment and this processed GP interacts with the endosomal receptor, NPC1, to facilitate fusion, leading to viral replication. Axl, a TAM receptor, is critical for EBOV entry into primary and immortalized keratinocytes and fibroblasts (6). However, the mechanisms by which EBOV reaches the skin in vivo remain unclear.

In this study, we used EBOV in nonhuman primates (NHPs) and mouse adapted-EBOV (ma-EBOV) and the low containment model virus rVSV/EBOV GP in mice to examine EBOV-GP-mediated trafficking to (inside out) and from (outside in) the skin. Our findings demonstrate involvement of skin endothelial, myeloid, and epithelial cells in EBOV infection of these animal models. In the outside in studies, we find a critical role for epidermal infection prior to subsequent systemic infection. Further, we characterize the inflammatory environment during the infection and identify PS receptors that restrict skin infection. Together, these results suggest that virus routinely traffics and infects skin during EBOV pathogenesis.

## MATERIALS AND METHODS

### Generation and titering of virus stocks

EBOV kikwit-9510621 (P2) was obtained from Dr. Tom Ksiazek (at NIAID’s WRCEVA at UTMB’s Health Galveston National Laboratory) in 2012 and propagated at Texas Biomedical Research Institute. The stock virus was passaged for a third time in Vero E6 cells and had a titer of 2.1 x 10^5^ PFU/mL as determined by an infectious plaque assay on vero E6 cells. The EBOV exposure stock has been confirmed to be wild-type Ebola virus by deep sequencing.

ma-EBOV (Ebola virus, Mayinga, Mouse Adapted EZ-76) virus was propagated on Vero-E6 cells (ATCC #CRL-1586) in the presence of 2% FBS. When CPE was evident, the culture supernatant was collected, cell debris pelleted by centrifugation at 1000xg for 10 minutes and the clarified supernatant containing virus was stored frozen in 1 mL aliquots at −80°C. Virus titer was determined as described in a previous report (8). Briefly, Vero E6 cells are infected with ma-EBOV for 1 h and then overlayed with methylcellulose. After 3 days, the overlay was removed, cells were fixed in formalin and then stained with a virus-specific antibody. Foci of infection were counted and used to back-calculate the virus titer in focus forming units (FFU) per mL.

rVSV/EBOV GP *in vivo* viral stocks were generated and prepared by infecting Vero E6 cells with a low MOI (∼0.001). Supernatants were collected at 24 and 48 hours post infection when cytopathic effect was evident. Viral supernatants were filtered through 0.45 μM filters and concentrated by layering virus over a 25% sucrose/DPBS cushion and ultracentrifuging for 2 hours at 120,000g. Viral titers were determined through TCID_50_ assay where serial dilutions of rVSV/EBOV GP are added to wells in eight replicates per dilution containing 10,000 Vero E6 cells per well in a 96-well plate. Virus is incubated for five days and the TCID_50_ was determined on Vero E6 cells using the Reed and Muench method and written in infectious units (IU) per mL (9). For each stock of virus used, serial dilutions of rVSV/EBOV GP were injected intraperitoneally (ip) or painted on shaved, abraded skin of *Ifnar ^−/−^* mice to determine viral titers used in the studies. Weight loss, morbidity, and mortality were measured to assess the lethal dose (LD) of each stock generated per route of infection.

### In vivo studies

#### Ethics statements

Animal research at the Texas Biomedical Research Institute was conducted under an Institutional Animal Care and Use Committee (IACUC)-approved protocol (IACUC number 1482MM) in compliance with the Animal Welfare Act, Public Health Service (PHS) policy, and all applicable federal regulations governing animal experiments. NHP time course study was conducted under IACUC-approved protocol number 1604MM and approved by study veterinarians at Texas Biomedical Research Institute (10). The Texas Biomedical Research Institute adheres to the Guide for the Care and Use of Laboratory Animals published by the National Research Council and is accredited by the Association for Assessment and Accreditation of Laboratory Animal Care. All euthanasia procedures were approved by study veterinarians and performed in accordance with the recommended method of the Panel on Euthanasia of the American Veterinary Medical Association. Established euthanasia criteria were strictly followed to minimize pain and distress.

BSL4 ma-EBOV murine studies were performed in accordance with protocol # 201900062, that was reviewed and approved by the IACUC at Boston University. BSL4 work was conducted in accordance with all institutional, local, state, and federal regulations and guidelines for BSL4 containment work.

Animal research and breeding performed at the University of Iowa (UI) were conducted in accordance with the Animal Welfare Act and the recommendations in the Guide for the Care and Use of Laboratory Animals of the National Institutes of Health (UI Institutional Assurance Number: #A3021-01). All animal procedures were designed to minimize animal discomfort and were approved by the UI IACUC. The BSL2 rVSV/EBOV GP *in vivo* mouse study was performed in accordance with the IACUC guidelines (protocols #4021280 Filovirus glycoprotein/cellular protein interactions and virus glycoprotein/cellular protein interactions).

### Non human primate studies

Three independent experiments were used as sources of material from EBOV-infected or mock-infected rhesus macaques.

#### EBOV *in vivo* NHP time course infections

In the first study, NHP RNA samples were received from EBOV-infected NHPs euthanized over the course of infection as previously described (10). Briefly, 12 male and 12 female rhesus macaques aged 3 to 7 years old were used in this study. 20 NHPs were infected with 1000 PFU of EBOV (Kikwit) and 4 NHPs were mock infected with sterile PBS. All animals received an intramuscular (im) injection and euthanized on scheduled endpoints at 3-, 4-, 5-, and 6-days post infection or until endpoint was reached at 7- or 9-days post infection. Tissue from the site of injection was placed in TRIzol for viral load analysis. Additional details from this study have been previously described (10).

#### EBOV *in vivo* NHP time course infections

For the second study, material from two animals used as treatment controls in a study evaluating antivirals for EBOV. Each received saline only as treatment. NHPs were acclimated for seven days in the ABSL4 prior to infection. On day of exposure, Day 0, NHPs received an im inoculation of 1,000 PFU of (P3) Zaire EBOV strain Kikwit 95. Animals were observed at least twice daily for morbidity and mortality until endpoint was reached (day 6 and 8). Blood was collected on 0- and 5-days post exposure and upon death for determination of coagulation parameters, hematology, clinical chemistry, and viral load. Weight and temperature data were collected during sedation time points. Beginning at 6 (+/- 1) hours post exposure, then again at 16, 28 (+/- 2 hours) and daily on days 2-8 post EBOV exposure (+/- 2 hours), NHPs received an im dose of vehicle. Uninfected mock NHPs were euthanized at 28-days post mock im exposure with sterile PBS. Post-euthanasia, a representative sample of skin, including the epidermis, dermis, and subcutis with underlying muscle, was collected with a scalpel from the site of inoculation and directly placed in the standard tissue cassette. The cassette with the tissue was placed in 10% buffered formalin for fixation for a minimum of 21 days. The sample was then paraffin embedded for immunostaining.

### Mouse studies

WT C57Bl/6 mice were purchased from Jackson Lab (Bar Harbor, ME). *Ifnar ^−/−^* C57BL/6J mice were a kind gift from Dr. John Harty (University of Iowa, Iowa City, IA, USA) and bred in house. C57BL/6 *Tim-1 ^−/−^ Ifnar ^−/−^* and *Tim-4 ^−/−^ Ifnar ^−/−^* have been previously described (11, 12) and were bred at the University of Iowa. C57BL/6 *Axl ^−/−^*were obtained from Dr. Rolf Brekken (Univ. Texas, Southwestern, Houston, TX) and crossed onto a C57BL/6 *Ifnar ^−/−^* background generating C57BL/6 *Axl ^−/−^ Ifnar ^−/−^*. C56BL/6 *Tim-1 ^−/−^ | Axl ^−/−^ Ifnar ^−/−^* were generated by crossing C57BL/6 *Tim-1 ^−/−^ Ifnar ^−/−^* with C57BL/6 *Axl ^−/−^ Ifnar ^−/−^* and bred at the University of Iowa. C57BL/6 *Tim-4^−/−^ | Axl ^−/−^ Ifnar ^−/−^* were generated by crossing C57BL/6 *Tim-4 ^−/−^ Ifnar ^−/−^* and C57BL/6 *Axl ^−/−^ Ifnar ^−/−^*and bred at the University of Iowa. Mouse genotypes were verified using primers described in Table 1 and PCR conditions previously described (11). Briefly, as approved and described by the UI IACUC policies and guidelines, a small <5 mm of the mouse tail is snipped and processed using the Wizard Genomic DNA Purification Kit by Promega as described by the manufacturer’s instructions. DNA is quantified by nanodrop. Per each reaction, 100 ng of genomic DNA, primers, and the New England Biolabs One*Taq* 2x Master Mix are mixed and a touchdown PCR protocol is used as recommended by Jackson labs.

**Table 1.**
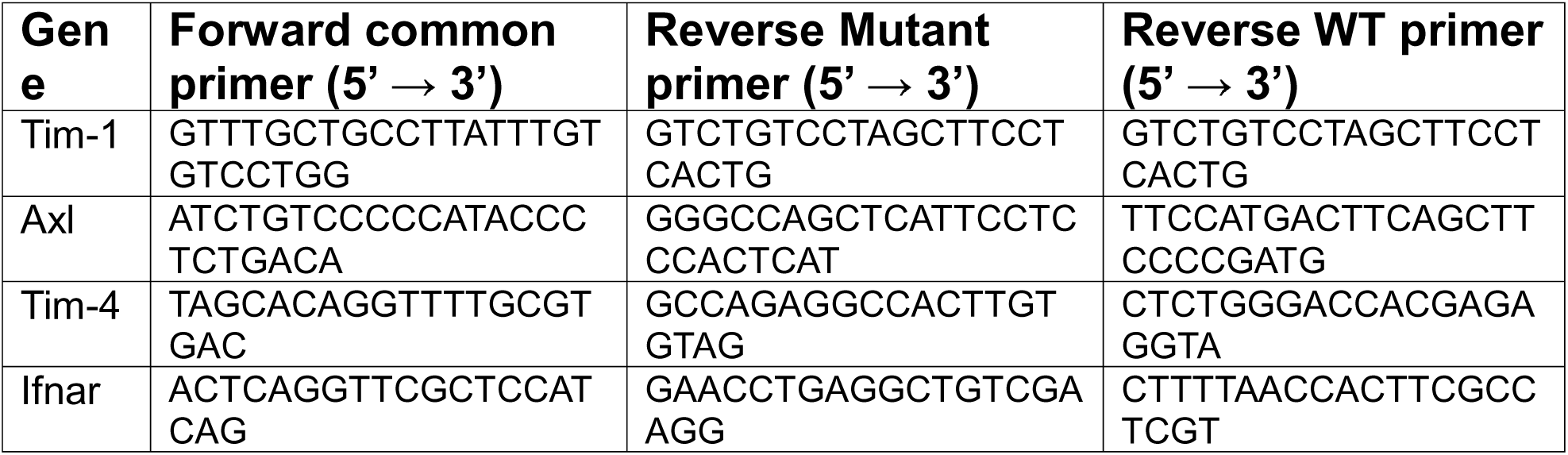
Mouse genotyping primers used in this study.

#### ma-EBOV *in vivo* infections

6-week old, female C57BL/6 N mice were infected ip with 1000 FFU of ma-EBOV (13). Animals (n=10) were euthanized every 24 hours after infection for 5 days. Tissues collected at specified time points were liver, subcutaneous fat associated with the belly, visceral fat in the belly region, belly skin (adjacent to injection site), back skin (distal to site of injection), and inner thigh skin (distal to site of injection). Tissue was either processed in TRIzol Reagent to measure transcriptional changes or fixed in formalin for 3 days and then paraffin embedded for immunostaining studies, described in detail below.

#### rVSV/EBOV GP *in vivo* infections

To investigate EBOV-GP mediated infection to distal skin, 6-10 week old, male mice on an *Ifnar ^−/−^* C57BL/6J background were shaved on the right back flank and subsequently infected ip with 500 IU of rVSV/EBOV GP (∼LD_100_) within 3-5 days. Specific mouse genotypes, time points, and number of mice can be found in the figure legends.

Focal infection was specifically visualized by shaving a much larger region of back skin in 6-10 week old, *Ifnar ^−/−^* C57BL/6J male mice. Following a five-day waiting period, mice were ip infected with a lethal dose of rVSV/EBOV GP. Mice were euthanized when endpoint criteria was met. Back tissue was removed, placed on a microscope slide, and visualized on an epifluorescence microscope. Areas of focal infection were marked by sharpie and captured using a camera. 3 mm dermal biopsy punches were taken from several areas where GFP was and was not evident. Individual biopsies were weighed and placed into Eppendorf tubes with 0.5 mL DPBS, homogenized using 1.5 mL Eppendorf pestles, and stored in −80°C.

EBOV-GP mediated viral dissemination from skin was also investigated. 6-10 week old, male mice on an *Ifnar ^−/−^* C57BL/6J background mice were shaved on the right back flank. 5 days after shaving, mice skin was gently abraded with an emery board until the mouse back was shiny (abraded). Another set of mice did not undergo abrasion (painted). 10 μL aliquot of concentrated virus was placed on the surface of the skin and mice were monitored until virus dried on the skin. Weight loss, morbidity, and mortality was observed throughout the course of infection. Surviving mice were kept for 21-days and then bled using a goldenrod animal lancet to poke the submandibular vein. Blood was collected in a microtainer blood collection tube (BD SST, Gold) and spun for 10,000 g for 10 minutes. Serum was collected, placed in an Eppendorf tube and kept at 4°C to later measure EBOV-GP specific serum IgG antibody titers as described below.

Mice were euthanized from 6 hrs to 3 days post infection and the spleen, liver, back were collected. In some experiments, the serum, testes, cheek and belly were also taken. Serum was collected by bleeding mice using a goldenrod animal lancet to poke the submandibular vein. Blood was collected in microtainer blood collection tube (BD SST, Gold) and spun for 10,000 g for 10 minutes. Serum was collected, placed in an Eppendorf tube, and kept at −80°C for viral titering as described below. Portions of the back and cheek were also fixed in formalin for 24 hours and later paraffin-embedded for immunostaining. All tissues were weighed prior to homogenization in 0.5 mL DPBS using 1.5 mL Eppendorf pestles.

Specific viral kinetics in epidermal and dermal tissue was measured in enzymatically dissociated back tissue. 5, 4mm punch biopsies were taken from the shaved back and submerged into 5 mg/mL of dispase ii solution (DPBS) for 2 hours at 37°C. The epidermis was separated from the dermis using forceps. The epidermis and dermis per mouse were weighed and homogenized in 0.5 mL DPBS. All tissue was placed in −80°C for future viral titering.

### NHP explant studies

#### NHPs explant infection

1 cm² NHP skin explants were generated from rhesus macaque skin samples (n=2 macaques) and placed on semipermeable transwells (2.3 cm diameter, 3.0 µM pore size; Falcon 353091). Explants were kept up to 12 days post infection(dpi) and maintained on 1.5 mL of MEM media supplemented with 10% FBS, 1% PS, fungizone and 0.2 µg/ml of type I interferon (IFN) inhibitor B18R. Explants were infected with 1.5×10^7^ IU of rVSV/EBOV GP in the basal media. In another set of experiments, 1.5×10^7^ IU of rVSV/EBOV GP was placed on the apical epidermal surface in 1.5 μL and allowed to dry in the biological safety cabinet. Following overnight incubation, explants were washed twice with DPBS and media was replaced. Supernatants were collected to measure viral infection by TCID_50_ and media was replaced every other day. Tissue was fixed on day 8 post infection in 4% paraformaldehyde overnight at 4°C for immunostaining.

### Assays detecting virus infection and replication

#### TCID_50_ studies

Vero E6 cells were plated at 10,000 cells per well in a 96-well format. The following day, homogenized tissues and serum were allowed to thaw to room temperature and filtered through 0.45 μM filters. Serial dilutions of infected tissues were plated in 6 technical replicates. Virus is incubated for five days and the TCID_50_ was determined on Vero E6 cells using the Reed and Muench method and written as TCID_50_ IU per gram or mL. All tissues only underwent one freeze-thaw cycle.

#### RT-qPCR

ma-EBOV tissue was homogenized in TRIzol LS using a Qiagen TissueLyzer with a 5 mm stainless steel bead according to the manufacturer’s directions, running 2 cycles, 2 min each at 30Hz. Samples were then clarified by centrifugation, frozen at −80°C and shipped to the University of Iowa after being certified to be inactivated. Samples were further processed to extract RNA by manufacturer’s instructions. Total RNA was quantified by nanodrop in order to calculate 1 μg of RNA. cDNA was generated using the High-Capacity cDNA Reverse transcriptase kit from the 1 μg of RNA as described by the manufacturer’s instructions. Using PowerUP SYBR Green master mix, quantitative PCR (qPCR) was performed to calculate the ΔCT values in relation to the house keeping gene. Specific primers can be found in Table 2.

**Table 2.**
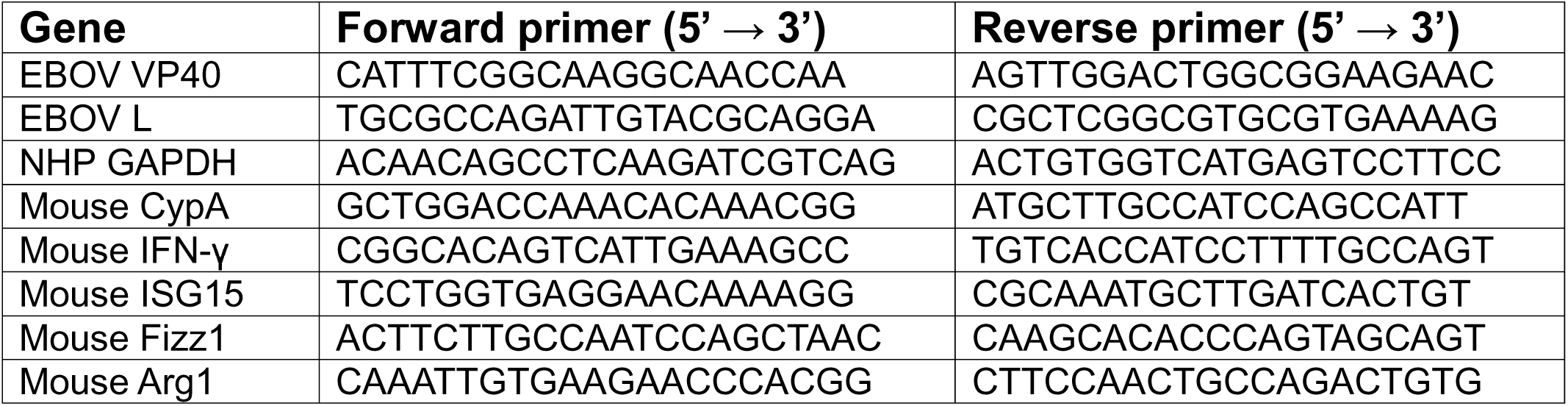
Oligonucleotides primers used in this study.

### Anti-EBOV GP IgG ELISAs

Detection of anti-EBOV GP antibodies was performed by coating optical MicroTiter plates (Immulon®) overnight with soluble EBOV GP (Kikwit-95, Acro Biosystems) (50 µL/well at 1 µg/ml). A standard curve of mouse immunoglobulin was generated and plated duplicate in non-coated wells. Wells were washed four times with ELISA Wash Buffer (1X PBS + 0.015% Tween 20), blocked for 2 hours at 25°C with ELISA Blocking Buffer (1X PBS + 2% BSA; 200 µL/well) and incubated overnight at 4°C with three serum dilutions (1:1000, 1:10,000, and 1:100,000) in duplicate obtained from mice surviving rVSV/EBOV GP skin infections. Following incubation, wells were washed as described above and incubated with horseradish peroxidase (HRP)-conjugated anti-mouse IgG (in 50 µL ELISA blocking buffer; 1:500 dilution of antibody) for 1 hour at 25°C. Wells were washed as described and incubated with HRP substrate according to the manufacturer’s protocol (TMB Substrate). The reaction was stopped with 50 µL 2M H2SO4 and absorbance at 450 nm was measured on a Tecan Plate Reader (Life Sciences). Total concentration of anti-EBOV GP antibodies was quantified by comparison to the standard curve.

### H&E staining of animal tissues

#### Hematoxylin and Eosin (H&E) staining

Fixed tissues were delivered to the Comparative Pathology Core (University of Iowa) and consistently processed (dehydrated through a progressive series of alcohols and xylene), paraffin embedded into blocks, and sectioned (∼4-5 µm) onto glass slides. The tissues were routinely stained with hematoxylin and eosin (HE) stain. Tissues were examined by a boarded veterinary pathologist.

### Primary murine fibroblast studies

#### Generation of primary murine fibroblasts

In sterile tissue culture hoods, mouse skin was dissected and were washed in 70% ethanol and povidone-iodine solutions followed by several DPBS washes. Small sections of mouse skin tissue was embedded into 10 cm^2^ tissue culture treated plates with a scalpel. Tissue was maintained in DMEM with 10% FBS and 1% PS and replaced weekly. Within 2-3 weeks, fibroblasts grew out from the skin sections creating a monolayer. Cells were passaged one or two times before infection experiments.

#### Primary murine fibroblasts infection

All primary fibroblasts were cultured in triplicate in a 48-well plate at ∼100,000 cells per well in a volume of 400 μL of DMEM with 10% FBS and 1% PS. Fibroblasts were infected for 48 hours at an MOI of 10 with rVSV/EBOV GP. Cells were detached with 0.25% trypsin/EDTA, neutralized with equivalent volume of newborn calf serum, centrifuged, and resuspended in DPBS with 2% calf serum with azide. Cells were analyzed by flow cytometry on Beckman CytoFLEX to quantify the frequency of GFP^+^ cells.

### Statistical analysis

Statistical analysis was completed in Prism. Graphs on a log scale had statistics performed on log-transformed values. Statistical tests included both Student’s *t* test and a one-way analysis of variance (ANOVA). Flow cytometry analysis was done in FlowJo. More specific details regarding statistical tests and exact *n* per group can be found in the corresponding figure legends.

## RESULTS

### EBOV traffics to NHP skin at late times post infection

Non-human primates (NHPs) are deemed to be the gold standard experimental animal for modeling EBOV infection, closely mirroring human disease (10, 14). Prior studies report viral antigen-positive cells in both human and NHP skin at late infection times (5, 15). Low levels of infectious virus persists on macaque skin for up to three days postmortem, supporting anecdotal reports of human skin transmission during the largest EBOV outbreak in 2013 (4, 5, 16). This suggests EBOV traffics to the skin late in infection; however, the kinetics and specific cell types involved remain unclear. To investigate this possibility, we analyzed NHP skin samples from three different studies. All of the samples that we evaluated were from animals that served as controls in studies, receiving EBOV without antiviral or vaccine treatment.

We evaluated the quantity of viral RNA in skin from EBOV-infected NHPs. Macaques were infected im with 1000 PFU of EBOV (Kitwik) and euthanized on specific days (3–9) over the course of infection. Details of clinical findings and virus loads in other tissues aside from the skin were described previously (10).Viral RNA levels in skin at the injection site increased over time (**Fig. 1A**), suggesting there is ongoing replication in the skin that peaks at late stages when animals were near death.

**Figure 1.**
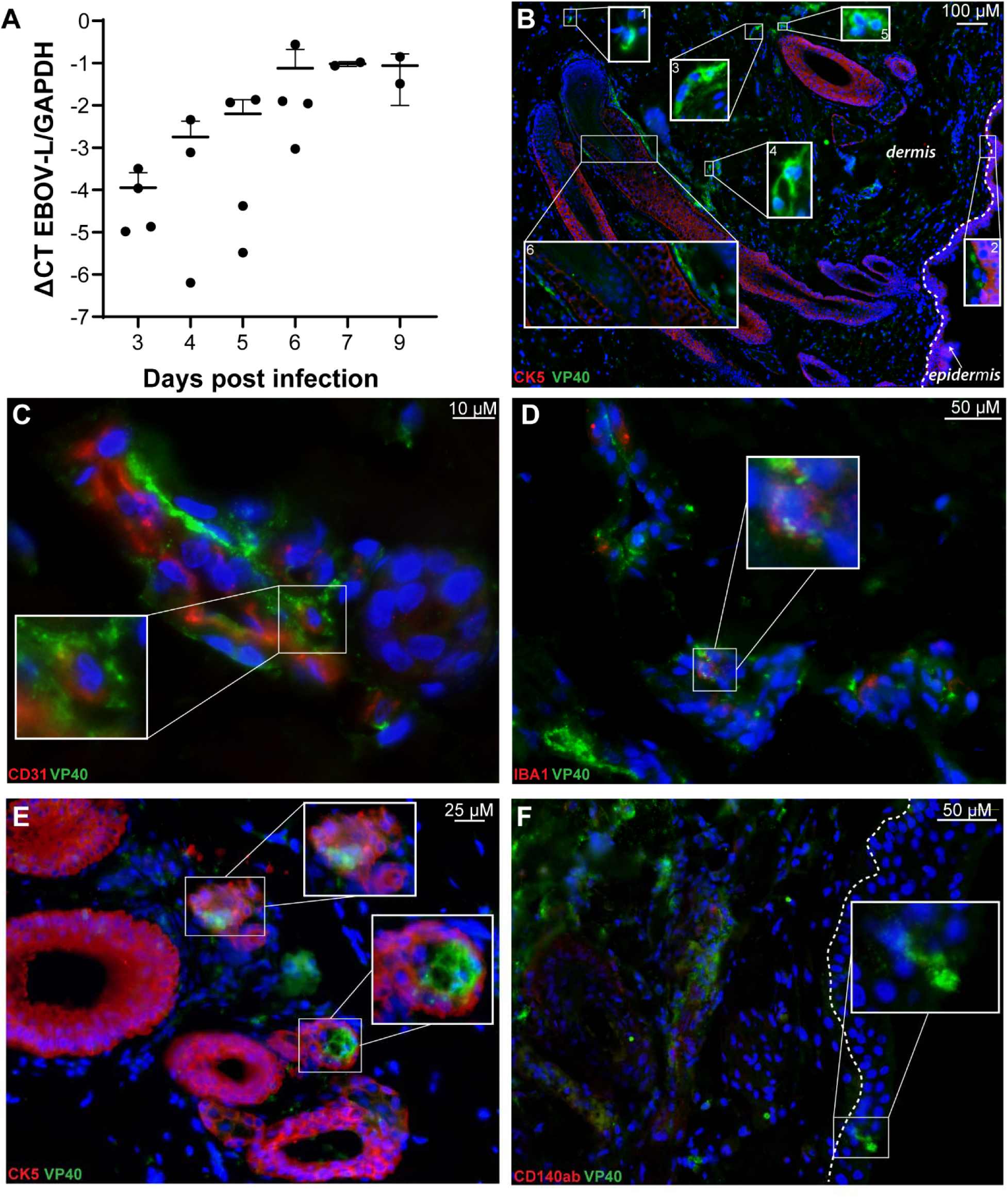
EBOV replicates and traffics to NHP skin at late times of systemic infection. NHPs were infected im with either 1000 FFU (**A**) or 100 FFU (**B-F**) of EBOV (Kikwit). NHPs were euthanized on specified days (**A**) or when endpoint criteria was met (**A-F**). EBOV L RNA was normalized in skin tissue at the site of injection compared to housekeeping gene, GAPDH (**A**). Data is expressed by ΔCT values and plotted on a log_10_ scale as means ±SD. No statistical significance was observed by ordinary one-way ANOVA (**A**). FFPE sections were taken contralateral (**B-E**) to or at (**F**) the site of injection at either 6 (**B**) or 7 (**C-F**) days post infection. Sections were mounted in DAPI (blue) and stained with virus (EBOV GP or VP40 in green) and cell markers CK5, CD140ab, CD31, and IBA1 in red. Entire images were enhanced by increasing brightness and decreasing contrast for visualization.

While formalin fixed, paraffin-embedded (FFPE) NHP skin tissues from this study were not available, FFPE NHP skin tissue was available from another EBOV (Kikwit) study in which male and female rhesus macaques where inoculated im with 100 PFU of EBOV (Kikwit) followed by termination at day 6 or 7. FFPE skin tissues were sectioned and immunostained. Sections from both skin adjacent to and contralateral from the injection site were immunostained for EBOV viral antigen (VP40 or GP) and cell lineage markers to define infected dermal cell types. Cell markers included CK5 (keratinocytes/epithelial cells), CD140ab (fibroblasts/smooth muscle/pericytes), CD31 (endothelial cells), and IBA1 (macrophages/microglia). Staining protocols and antibody details are provided in **Table 3**.

**Table 3.** Antibodies used in immunostaining studies.

| **Antibody description** | **Vendor/Item ID** | **Dilution** | **Incubation conditions** |
| --- | --- | --- | --- |
| Human anti-Ebola Virus GP mAb (KZ52) | IBT Bioservices / 0260-001 | 1:100 | Overnight / 4°C |
| Anti-MARV GP chimeric monoclonal antibody | IBT Bioservices / 02101-016 | 1:100 | Overnight / 4°C |
| ASGR1 Polyclonal antibody | Proteintech / 11739-1-AP | 1:50 | Overnight / 4°C |
| CD31 Polyclonal Antibody | Invitrogen / PA5-16301 | 1:100 | 2hrs / RT |
| Cytokeratin 5 Polyclonal Antibody | Invitrogen / PA5-29670 | 1:200 | 2hrs / RT |
| Anti-IBA1 Polyclonal Antibody | Fujifilm / 019-19741 | 1:300 | 2hrs / RT |
| Goat anti-Rabbit IgG, Alexa Fluor 488 | Invitrogen / A-11008 | 1:800 | 1hr / RT |
| Goat anti-Human IgG, Alexa Fluor 647 | Jaksonimmuno/ 109-605-003 | 1:1000 | 1hr / RT |
| Goat anti-Mouse IgG, Alexa Fluor 488 | Invitrogen / A-11029 | 1:800 | 1hr / RT |
| Goat anti-Rabbit IgG, Alexa Fluor Plus 555 | Invitrogen / A-48283 | 1:800 | 1hr / RT |

Substantial viral antigen positive cells were detected in contralateral skin by day 6 (**Fig. 1B**). Inserts highlight widespread clusters of infection, with VP40 antigen present from the deep dermis (**Fig. 1B, insert 1**) to the epidermal-dermal junction (**Fig. 1B, insert 2**). EBOV-positive cells included elongated fibroblastic- or pericyte-like cells (**Fig. 1B, inserts 3 and 4**) and rounded cells consistent with myeloid cells morphology (**Fig. 1B, insert 5**). This indicates a diversity of cell types supporting EBOV infection within the skin. EBOV antigen was frequently found in cells adjacent to but not within CK5+ hair follicles (**Fig. 1B, insert 6**), suggesting potential infection of nearby mesenchymal cells involved in follicle maintenance (17). Specificity of immunostaining was demonstrated by sections stained with secondary antibody only and the absence of EBOV antigen staining in uninfected tissues (**Fig. S1A-B**). Since EBOV causes hemorrhagic fever, and NHPs exhibit its hallmarks, we examined viral colocalization with dermal vasculature. EBOV antigen was present in CD31-positive blood vessels (**Fig. 1C, inset**), suggesting endothelial cells support dermal infection. As expected, viral antigen also colocalized with the macrophage marker IBA1 within the dermis (**Fig. 1D, inset**). This is consistent with myeloid cells being early and sustained EBOV targets. Hair follicles harbored vast viral antigen within structures consistent with sebaceous glands, but only occasionally colocalized with CK5-positive cells in hair follicles (**Fig. 1E, Fig. S1C-D**). Infection of the epidermis was present but infrequent (**Fig. 1F**). Low levels of EBOV infection of the epidermal layer was unexpected. Earlier studies demonstrated prominent epidermal infection at late stages in human skin explants and detectable infectious virus on the surface of postmortem NHP skin (5, 6).

To further assess NHP epidermal susceptibility to EBOV-GP-mediated, we used an explant model with 1 cm² rhesus skin sections at the air/liquid interface on semipermeable transwells. A biosafety level 2 (BSL2) EBOV model virus, rVSV/EBOV GP, recombinant vesicular stomatitis virus (rVSV) expressing EBOV GP and GFP, was used in the presence of a STAT1 inhibitor, B18R. rVSV/EBOV GP tropism in the presence of an interferon inhibitor closely mimics BSL4 EBOV tropism (6). Viral infectious titers in basal media increased over time (**Fig. S1E**), and viral antigen was detected in the dermis and epidermis. Viral antigen within the epidermis co-localized with CK5+ cells (**Fig. S1F**), indicating that NHP keratinocytes are permissive to rVSV/EBOV GP. Together, these findings characterize EBOV infection kinetics and the diverse infected cell populations within NHP skin.

### ma-EBOV traffics to mouse skin at late stages of infection

To further investigate EBOV trafficking to the skin, we used a mouse model, which allows for a higher sample size and more precise kinetic analysis. While mice do not fully recapitulate human filovirus disease, they provide valuable insights into trafficking of the virus over the course of infection and the cells targeted for infection (14, 18). Six-week-old female C57Bl/6 mice were infected ip with 1000 FFU of ma-EBOV that generally results in lethality between days 6–8 (18). Mice (n=10 per day) were euthanized on days 0-5 of infection, and tissues were analyzed via RT-qPCR and immunostaining.

The liver, a known EBOV target, served as a positive control. As expected, viral RNA levels increased significantly in a time-dependent manner, reaching statistical significance within one day and plateauing on days 3-5 (**Fig. 2A**). Similarly, visceral fat near the injection site showed increasing viral load over time (**Fig. 2B**). Thigh skin, distal to the injection site, showed a delayed but significant viral presence by day 5 (**Fig. 2C**). Other tissues that were assessed for viral load included skin at the site of infection (belly skin), subcutaneous fat lining the belly skin, and skin taken from the back of mice (**Fig. S2A-C**).

**Figure 2.**
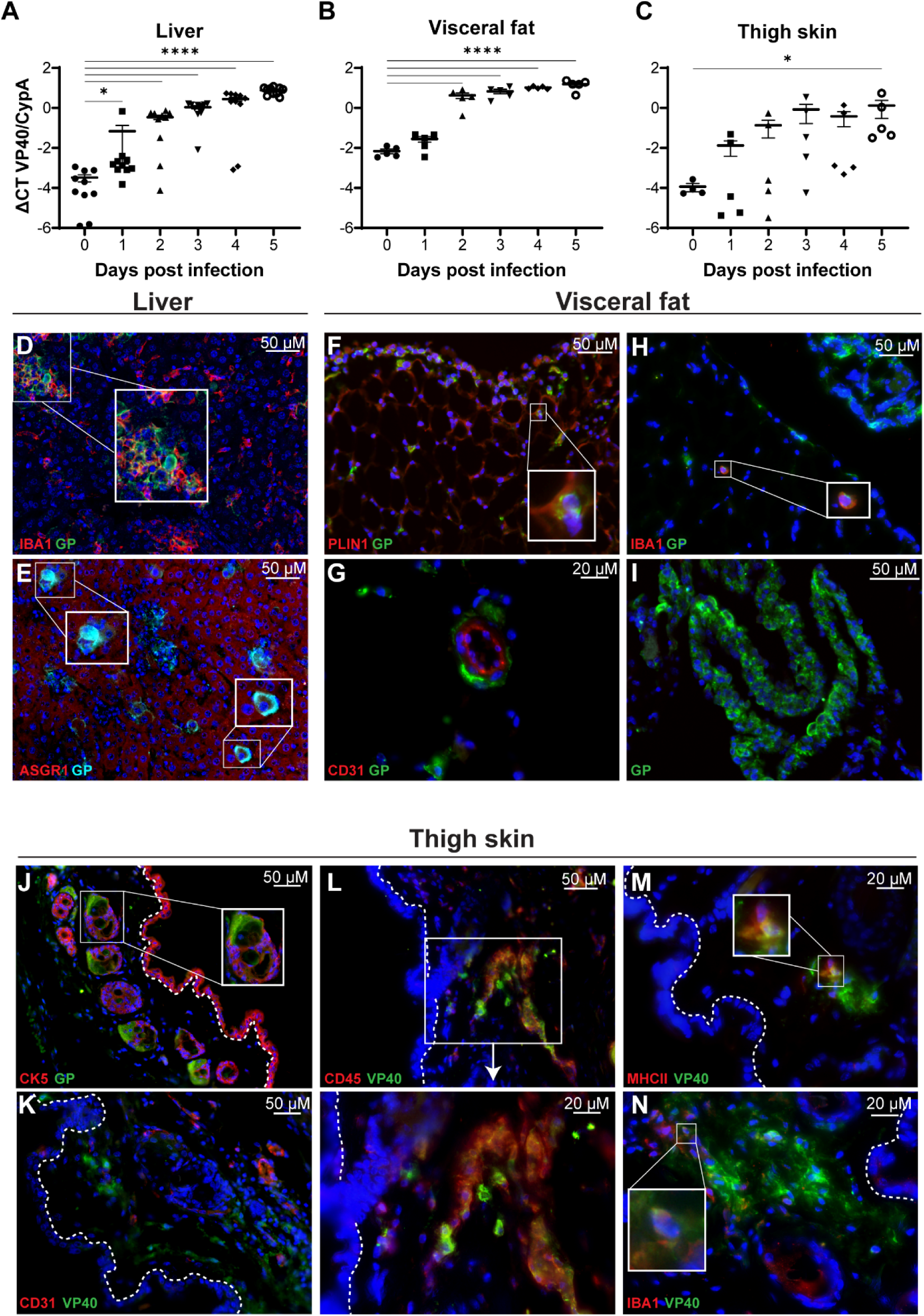
Adjacent and distal mouse tissues have distinct ma-EBOV kinetics and viral cell tropism. 6-8 week old female C57BL6/N mice were infected with 1000 FFU of ma-EBOV by intraperitoneal injection. N=10 mice were euthanized daily and liver, visceral fat, subcutaneous fat, belly skin (adjacent to site of injection), thigh skin (distal from site of injection), and back skin (distal from site of injection) tissues were collected. Tissues were either processed for RT-qPCR in TRIzol (**A-C**) or FFPE for immunostaining (**D-N**). **A-C**) Viral load was assessed by qRT-PCR in the liver (**A**), visceral fat (**B**), and thigh skin (**C**) over time. Viral load (VP40) was normalized to the house keeping gene cyclophilin-A (CypA). Data is expressed on a log_10_ scale as means ±SEM. Ordinary one-way ANOVA, *p<0.05 ****p<0.0001. **D-N**) FFPE sections were taken from 5 days post infection in the liver (**D,E**), visceral fat (**F-I**), and thigh skin (**J-N**) to stain for EBOV viral antigen (GP and VP40, green), cell lineage markers (IBA1, ASGR1, PLIN1, CD31, CK5, CD45, MHC II, red), and mounted in DAPI (blue). Insets demonstrate colocalization of cell lineage markers and viral antigen. Entire images were enhanced by increasing brightness and decreasing contrast for visualization.

To identify infected cell types, we performed immunostaining on tissue sections from late-stage infection. Cell specific markers included ASGR1 (hepatocytes), CK5 (keratinocytes/epithelial cells), PLIN1 (adipocytes), CD31 (endothelial cells), CD45 (immune cells), MHC II (antigen-presenting cells), and IBA1 (macrophages/microglia). Controls confirmed specific staining (**Fig. S2D–E**).

Consistent with previous reports, we detected EBOV antigen in liver macrophages (Kupffer cells) and hepatocytes at day 5 post infection (**Fig. 2D–E**). Focal areas of infection were also observed (**Fig. 2D, inset**), aligning with prior ma-EBOV studies (19). In visceral fat that was adjacent to the peritoneal compartment (the site of injection), viral antigen colocalized with PLIN1+ cells, confirming adipocyte infection in vivo (14) (**Fig. 2F-I**). While endothelial cells in the visceral fat were not found colocalize with viral antigen, infected cells were observed around blood vessels (**Fig. 2G**). This aligns with previous findings that endothelial infection is rare outside the liver and lymphoid tissues in ma-EBOV-infected mice (20). Although macrophages are key EBOV targets, only occasional IBA1+/EBOV antigen+ cells were detected in visceral fat (**Fig. 2H**). The most prominent viral antigen-positive cells were found in peritoneal ligaments that are fold of serous membranes connecting visceral organs (**Fig. 2I, Fig. S2G-H, red arrows**). H&E stained sections of visceral ligaments had notable areas of mononuclear inflammation in the visceral ligaments (**Fig. S2G-H, red arrows)** and parts of the lining of the visceral peritoneum (**Fig. S2G-H, black arrows**).

In skin that was distal to the site of infection sites, such as thigh skin, abundant clusters of viral antigen-positive cells were observed near the epidermal-dermal interface at day 5 (**Fig. 2J–N**), similar to NHP skin (**Fig. 1**). EBOV antigen localized along hair follicles and possibly sebaceous glands (**Fig. 2J, Fig. S2F**). As seen in visceral fat, viral antigen did not colocalize with endothelial cells (**Fig. 2K**). By day 4, small clusters of infected CD45-positive immune cells appeared in the dermis (**Fig. S2I-J**), increasing by day 5 (**Fig. 2L**). Some infected cells expressed MHC II, indicating the presence of antigen-presenting cells in the dermis (**Figure 2M**). Surprisingly, while macrophages were abundant in these foci, few were EBOV-positive (**Figure 2N, inset**). Together, these findings suggest that at late stages of infection ma-EBOV traffics to skin, infecting multiple cell types, including adipocytes, fibroblasts, mesothelial cells, and immune cells, while largely sparing endothelial cells.

### ma-EBOV induces distinct inflammatory transcription profiles in mouse tissues

EBOV upregulates proinflammatory cytokines in hosts despite encoding proteins that inhibit host antiviral responses (3, 20–30). We examined expression of inflammatory genes in ma-EBOV infected mice at day 5 post infection (**Fig. 3**). As expected, proinflammatory gene transcripts, ISG15 and IFN-γ, were elevated in the liver (**Fig. 3A, B**), while anti-inflammatory gene transcripts remained unchanged (**Fig. 3C, D**). A previous study had shown that adipocytes upregulate proinflammatory cytokine RNAs following EBOV and rVSV/EBOV GP infection (20). Consistent with this, proinflammatory gene, ISG15 and IFN-γ, transcripts increased in visceral fat (**Fig. 3E,F**). Interestingly, Fizz1 expression decreased while Arg1 increased, indicating a mixed inflammatory environment (**Fig. 3G,H**). In contrast, subcutaneous fat associated with the belly skin had elevated proinflammatory gene transcripts and no changes to the anti-inflammatory transcript levels (**Fig. S3A-D**). Belly skin also had a mixed inflammatory environment, potentially influenced by the proinflammatory profile of the subcutaneous fat (**Fig. S3E-H**). Distal skin tissue, thigh and back, however, had no significant changes in pro- or anti-inflammatory gene expression (**Fig. 3I-L, Fig. S3I-L**). This may be attributed to the extensive surface area of skin and the focal areas of infection (**Fig. 2J-N**).

**Figure 3.**
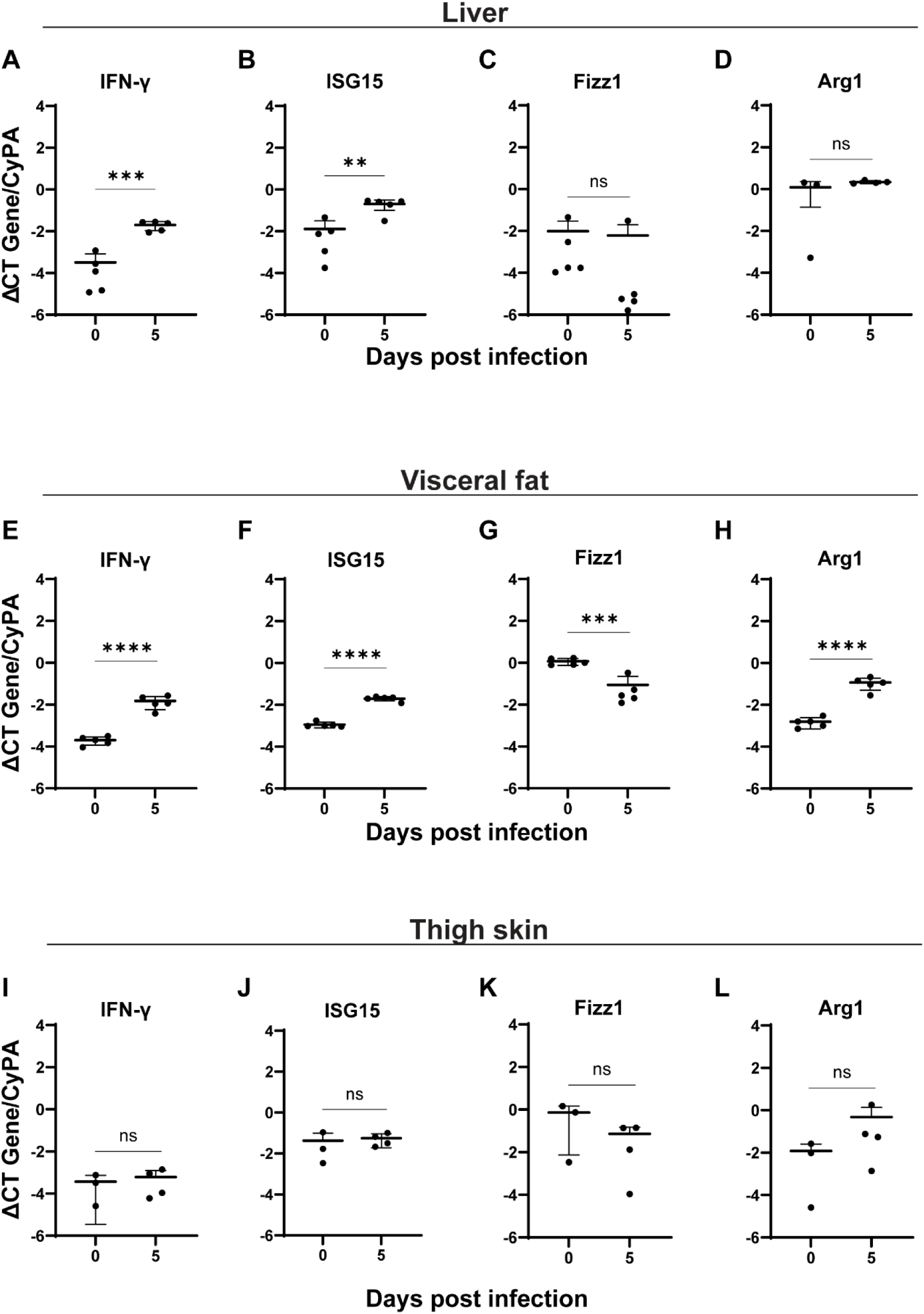
Inflammatory profiles of multiple mouse tissues infected with ma-EBOV. C57BL6/N mice were infected and tissues were collected and processed as described in Figure 2. Proinflammatory (IFN-γ and ISG15) and anti-inflammatory (Fizz1 and Arg1) gene transcripts were measured in the liver (**A-D**), visceral fat (**E-H**), and thigh skin (**I-L**). Values were measured in samples taken at 0 and 5 days post infection and normalized to the house keeping gene cyclophilin-A (CypA). Data is expressed on a log_10_ scale as means ±SD. Student’s t test, **p<0.01 ***p<0.001 ****p<0.0001.

### rVSV/EBOV GP infection models EBOV infection to mouse skin

To further investigate EBOV GP-specific tropism, we assessed rVSV/EBOV GP infection in C57BL/6 *interferon alpha-beta receptor knockout* (*Ifnar ^−/−^*) mice. rVSV/EBOV GP is lethal to these mice within 3–5 days post-infection, progressing more rapidly than EBOV (11, 31, 32). Viral titers were detectable in the spleen and liver within one day post-infection, reaching high levels by day three (**Fig. 4A**). Surprisingly, virus replication was limited in skin tissues near the injection site but reached high titers in distal back skin by day 2 (**Fig. 4A**).

**Figure 4.**
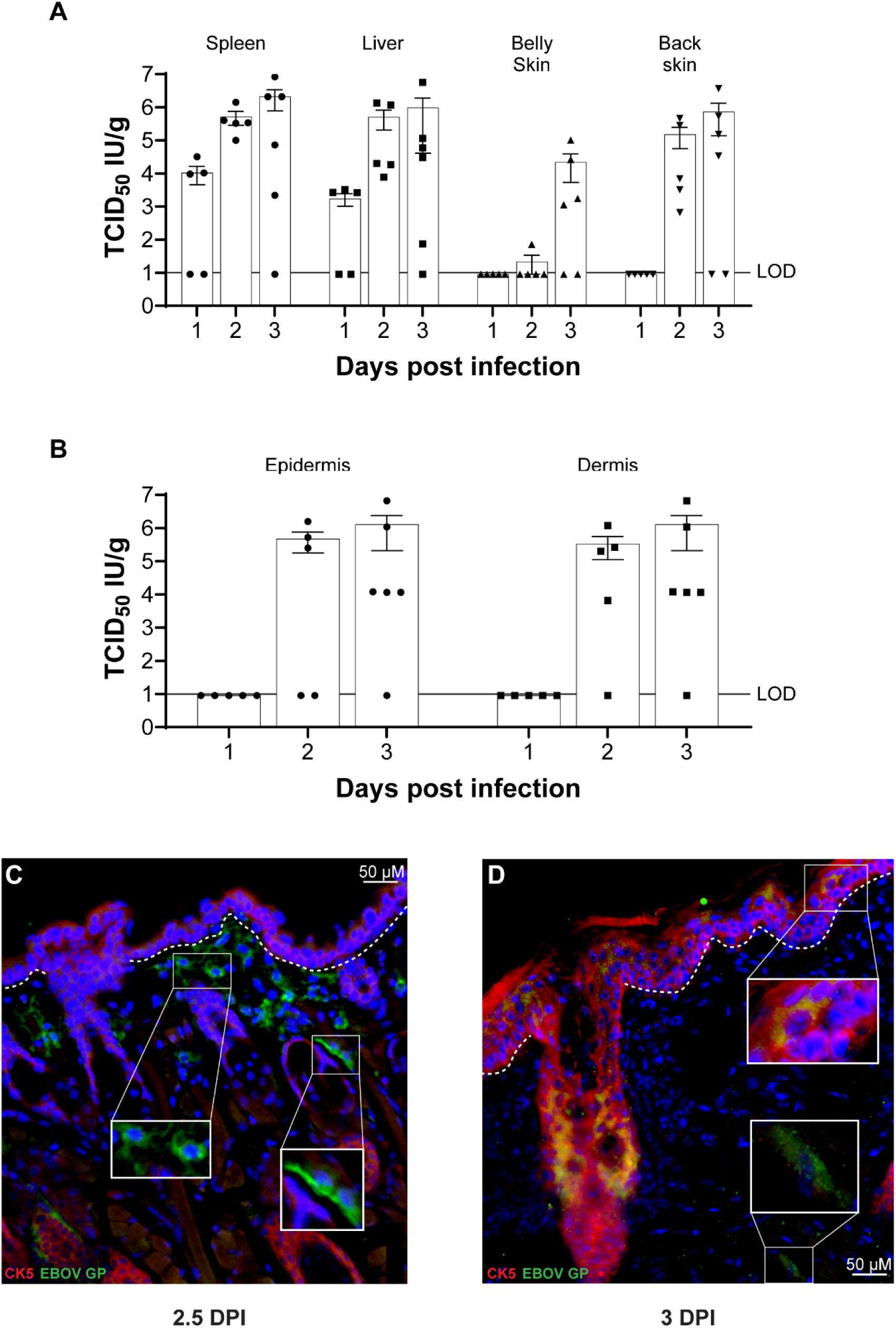
rVSV/EBOV GP mouse infections are similar to ma-EBOV. 6-8 week old male *ifnar ^−/−^* C57BL6/J mice were infected with 500 IU of rVSV/EBOV GP (n=6 per day). **A**) Infectious titers were measured in spleen, liver, belly skin, and back skin. Tissues were collected on 1, 2, and 3 days post infection, homogenized, and titered on Vero E6 cells. **B**) Back tissue was enzymatically dissociated to separate the epidermis and dermis, homogenized, and titered on Vero E6 cells. **C,D**) Mice were euthanized at 2.5 and 3 days post infection and skin was taken from the cheek for immunostaining. FFPE sections were cut and stained for viral antigen (EBOV GP, green) and keratinocyte marker (CK5+, red). **C**) Insets show infected CK5-negative cells that are round in shape (left) and elongated (right). **D**) Viral antigen was seen in CK5-positive epidermis (top inset) and CK5-negative cells in the dermis (bottom inset). Entire images were enhanced by increasing brightness and decreasing contrast for visualization.

To determine the contribution of the two layers of skin during viral dissemination, we enzymatically separated the epidermis and dermis from the skin of infected mice at 1-, 2-, and 3-days post infection. Viral titers were detectable by day two and peaked at day three, with both epidermal and dermal layers showing high viral loads (**Fig. 4B**). Immunostaining at 2.5 days post-infection revealed EBOV GP antigen in various dermal cell types, including immune and fibroblastic-like cells (**Fig. 4C**). By day three, viral antigen was also readily evident in keratinocytes in the epidermis and hair follicles (**Fig. 4D**). These findings suggest that rVSV/EBOV GP effectively models the viral trafficking and tropism observed in ma-EBOV infections *in vivo,* but more rapidly spreads systemically, resulting in skin infection at earlier times following inoculation. **rVSV/EBOV GP infection of skin can cause systemic infection** As our studies demonstrate that EBOV i.p. infection results in virus trafficking to distal skin locations (i.e., inside-out trafficking) and detectable virus replication in cells within the skin, we investigated whether skin could serve as an initial target of EBOV GP-mediated infection (outside-in trafficking) (6). Anecdotal reports suggest the possibility of person-to-person transmission through skin contact, prompting us to examine whether infection of skin can result in systemic infection (4, 16, 33). Using our low-containment virus model in NHP explants and *Ifnar ^-/-^* mice, we tested this hypothesis.

First, in our NHP skin explant transwell model, a small aliquot of concentrated rVSV/EBOV GP was applied to the skin surface and allowed to absorb. Infectious virus was monitored in basal media over an 8-day infection. As early as 2 days post-infection, infectious rVSV/EBOV GP was evident in basal media with significant increases by day 4 (**Fig. 5A**). At day 8, large foci of viral antigen were observed in the epidermis of FFPE sections (**Fig. 5B**), indicating that NHP keratinocytes are permissive to EBOV GP-mediated infection, similar to human keratinocytes (6).

**Figure 5.**
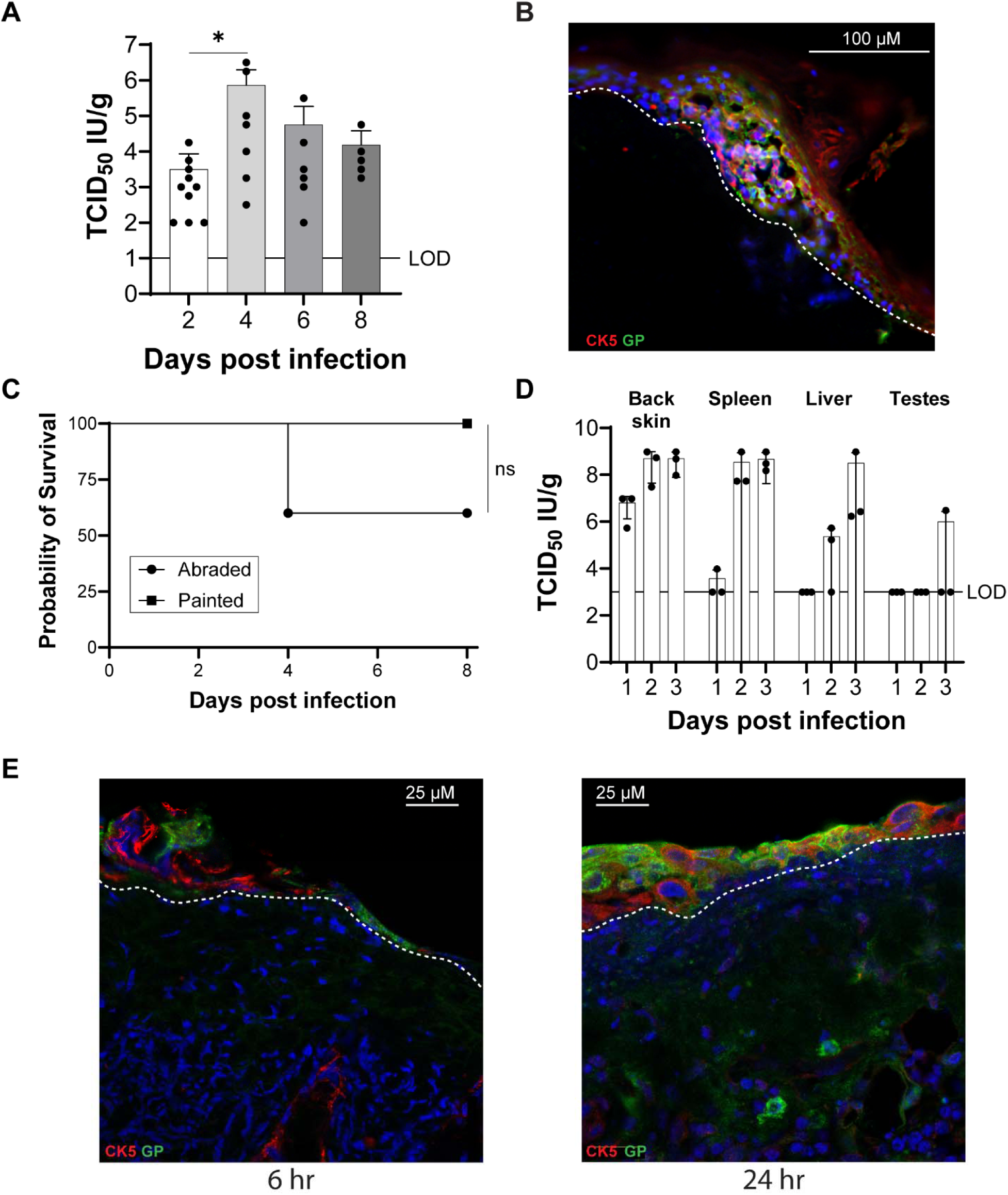
Direct infection of the skin epidermal surface results in viral dissemination and morbidity. **A,B**) 1 cm^2^ NHP skin explants from n=1 animal were generated from uninfected NHP skin. NHP skin explants were placed dermal side down on semipermeable transwell membrane and maintained at the air-liquid interface with media containing the STAT1 inhibitor, B18R. 1.5×10^7^ IU of rVSV/EBOV GP in 1.5 μL was placed on the apical epidermal surface. Following overnight infection, explants were washed twice with DPBS and media was changed every other day for 8 days. **A**) Production of infectious virus in the explants was measured in the basal supernatants over time on Vero cells. Shown are 1 independent experiments with data expressed on a log_10_ scale as means ±SD. Ordinary one-way ANOVA, *p<0.05. **B**) NHP skin explants were fixed on day 6 post infection in 4% PFA and stained for viral antigen (EBOV GP, green) and keratinocyte marker (CK5, red) mounted in DAPI (blue). The dotted line indicates the epidermal-dermal junction. **C**) 6-8 week old male *Ifnar ^−/−^* C57BL6/J mice (n=5) were shaved on the back right flank and allowed to rest for five days. 10 μL of 10^9^ IU of rVSV/EBOV GP was placed on the shaved skin either with (abraded) or without (painted) gentle abrasion. **C**) Survival was monitored and Log-rank test was performed (n=5 per group). **D-E**) 6-8 week old male *Ifnar ^−/−^* C57BL6/J mice (n=3) were shaved on the back right flank and allowed to rest for five days. 10 μL of 10^7^ IU of rVSV/EBOV GP was placed on the shaved abraded skin. Back skin, spleen, liver, and testes were collected on 1, 2, and 3 days post infection from abraded mice, homogenized, and titered on Vero E6 cells (n=3 per day). **E**) Abraded mice were euthanized at 6 (left) and 24 (right) hours post infection for immunostaining. FFPE sections were cut and stained for viral antigen (EBOV GP, green) and keratinocyte marker (CK5+, red). Entire images were enhanced by increasing brightness and decreasing contrast for visualization.

To assess in vivo dissemination, we compared two methods of skin infection: painting and abrasion. A patch of skin on the back of mice was shaved to remove the fur, rested for five days, then exposed to concentrated virus either with (abraded) or without (painted) gentle abrasion to the stratum corneum with an emery file (**Fig. 5C**). Infected mice were monitored over an 8 day period. ∼40% of the mice infected following abrasion succumbed to infection, suggesting that removal of the stratum corneum is necessary to facilitate direct skin infection, particularly with these high concentrations of virus (**Fig. 5C, Fig. S5A**). In the mice that received painted on virus, virus administration was much less virulent. Nonetheless, systemic dissemination occurred in abraded mice even at lower inoculum levels, as evidence by the presence of high concentrations of EBOV GP -specific IgG in serum (**Fig. S5B**).

Infectious virus titers were assessed in different tissues following virus administration to abraded skin. Skin taken from the infection site had high viral titers as early as 1 day post-infection, increasing by day 2 (**Fig. 5D**). Evidence of systemic infection was not observed until day 2 of infection with robust spleen and liver infection by day 2 and 3, respectively. Virus reached the testes in a single mouse by day 3. To visualize viral spread at the site of infection in the skin, mice were euthanized at 6- and 24-hours post-infection. At 6 hours, viral antigen was restricted to the epidermis in small foci (**Fig. 5E**). By 24 hours, infection had spread within the epidermis and reached immune-like cells in the deeper dermis (**Fig. 5F**). These findings demonstrate that rVSV/EBOV GP can infect and disseminate through the skin, supporting the possibility of skin-mediated EBOV transmission.

### Tim-1 mediates EBOV-GP viral trafficking to skin

A number of different cell surface receptors have been identified to enhance EBOV or rVSV/EBOV GP adherence and internalization into the endosomal compartment (11, 34–39). Phosphatidylserine (PS) receptors and C-type lectins have been most strongly implicated as important receptors in tissue culture studies. To date, only the PS receptor, TIM-1, has been shown to contribute to infection in vivo; at late times during infection, TIM-1 knock out (KO) mice have lower virus titers and reduced pathogenesis (11). To identify host cell surface receptors that are required for rVSV/EBOV GP to reach distal regions of the skin *in vivo*, we utilized *Ifnar ^−/−^* mice and crossed with whole-body knockouts of PS receptors, including *Tim-1, Tim-4, Axl,* and combinations thereof (*Tim-1^−/−^* | *Axl ^−/−^* and *Tim-4^−/−^* | *Axl ^−/−^*).

Male mice were infected ip with a lethal dose of rVSV/EBOV GP five days after shaving a patch of fur on the back. Mice were euthanized on day 3 of infection and infectious virus present in the spleen, liver, adjacent skin (belly), and distal skin (back) was determined (**Fig. 6A-E**). No differences in viral replication were observed in the spleen across PS receptor KO mice (**Figure 6A**), suggesting that systemic infection of the spleen is unaffected by these receptors at day 3 of infection. While titers in the liver tended to be more variable, there was no statistically significant differences between the strains (**Fig. 6B**).

**Figure 6.**
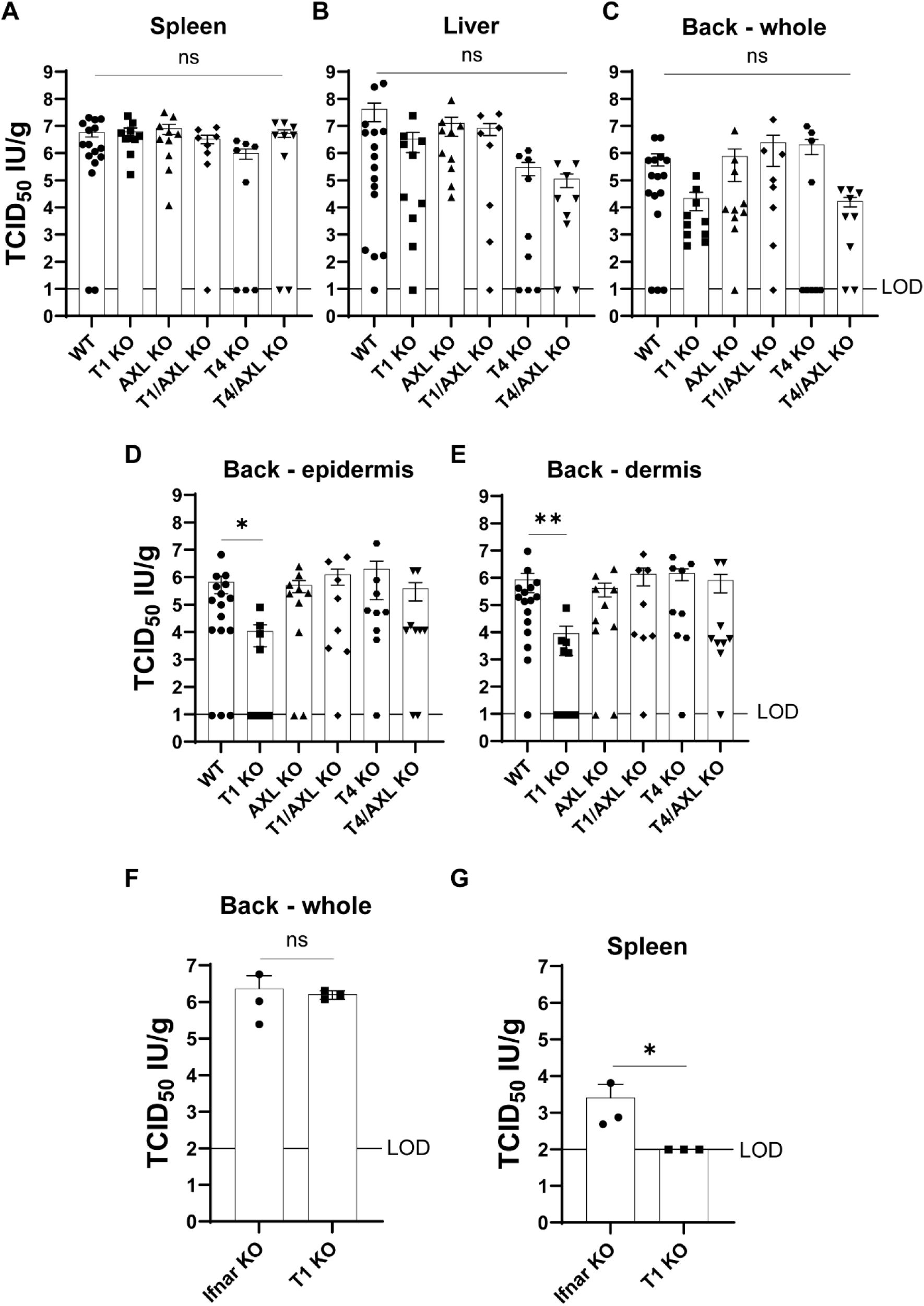
PS receptors mediate viral trafficking to and from skin. **A-E**) 6-10 week old *Tim-1 ^−/−^ Ifnar ^−/−^* (n=11), *Axl ^−/−^ Ifnar ^−/−^* (n=10), *Tim-4 ^−/−^ Ifnar ^−/−^*(n=11), *Tim-1^−/−^* | *Axl ^−/−^ Ifnar ^−/−^* (n=10), *Tim-4^−/−^* | *Axl ^−/−^ Ifnar ^−/−^*(n=10), and *Ifnar ^−/−^* (n=17) C57BL6/J mice were shaved on the back right flank and allowed to rest for five days. All mice were infected with 500 IU of rVSV/EBOV GP and euthanized at 3 days post infection. Spleen (**A**), liver (**B**), and intact back skin (**C**) tissue were collected, homogenized, and titered on Vero E6 cells. Back skin tissue was processed enzymatically dissociated to separate the epidermis (**D**) and dermis (**E**). The epidermis and dermis were homogenized and titered on Vero E6 cells. Ordinary one-way ANOVA, *p<0.05 **p<0.01. **F-G**) 6-10 week old *Tim-1 ^−/−^ Ifnar ^−/−^* (n=3) and *Ifnar ^−/−^* (n=3) C57BL6/J mice were shaved on the back right flank and allowed to rest for five days. 10 μL of 10^9^ IU of rVSV/EBOV GP was placed on the shaved skin following gentle abrasion. 24 hours after infection mice were euthanized and viral titers were measured in the intact back skin (**F**) and spleen (**G**). Statistics was determined by student’s T test on transformed values *p<0.05.

Infection in both belly and back skin showed high variability, likely due to focal infection patterns in skin tissue (**Fig. 6C, Fig. S6A**). Given the skin’s large surface area, it is not surprising that localized infections may have been missed. We also analyzed viral titers in skin epidermis and dermis (**Fig. 6D,E**). Surprisingly, *Axl ^−/−^* did not impact viral titers in either skin layer (**Figure 6D,E**), despite its established role in human dermal keratinocyte and fibroblast infection (6). However, primary mouse fibroblast infections confirmed *Axl*’s importance in vitro, similar to human fibroblasts (**Fig. S6B**). Notably, *Tim-1 ^−/−^* mice had significantly reduced viral titers in both the epidermis and dermis (**Fig. 6D,E**), suggesting an unexpected role for Tim-1 in promoting skin infection.

To assess if Tim-1 impacts in vivo dissemination from the skin, we infected *Ifnar ^−/−^* and *Tim-1 ^−/−^ Ifnar ^−/−^* through the back and euthanized 24 hours after infection. Following a five-day rest period after the back skin was shaved, a concentrated viral inoculum was placed on abraded skin. One day post infection, mice were euthanized and the back skin, spleen, serum, and liver were analyzed for infectious titers (**Fig. 6F,G**, and **Fig. S6C-F**). Tim-1 does not impact direct infection of the back skin (**Fig. 6F, Fig. S6C,D**). However, the *Tim-1 ^−/−^* had lower spleen and serum titers (**Fig. 6G, Fig. S6E**). Together, these data suggest Tim-1 plays a role in trafficking to and from the skin but does not impact direct infection of skin.

## DISCUSSION

Our findings demonstrate that EBOV GP-mediated viral trafficking to the skin occurs at later stages of infection. Skin is recognized as a critical site for EBOV replication and is used in clinical diagnostics (4). While our prior study explored direct EBOV infection of human skin explants (6), the in vivo kinetics and key cell types involved in viral trafficking in two important animal models had not been examined. Using two distinct animal models and three EBOV virus models, we identified both shared and unique features of skin infection. Viral replication in skin increased over time, underscoring its role as a key site of EBOV pathogenesis during systemic infection.

EBOV infects diverse skin cell types, and we consistently observed viral antigen in fibroblasts and immune cells, highlighting their importance in skin infection (6). Endothelial cell involvement was primarily observed in NHP skin tissue, though viral antigen was detected around vessels in dermis of murine tissue. Previous reports indicate minimal endothelial infection outside the liver and lymph nodes in ma-EBOV-infected mice, suggesting model-dependent differences in skin infection patterns (40). Epidermal infection was more pronounced in *Ifnar ^−/−^* rVSV/EBOV GP-infected mice, likely due to the rapid replication compared to EBOV and the absence of type I interferon signaling in these mice. The focal nature of infection in the skin may have also contributed to sporadic detection in ma-EBOV and rVSV/EBOV GP mouse models, as viral antigen may be present in unexamined regions. It is also likely that epidermal infection may be less apparent than dermal infection in animal models. In all animal models, EBOV antigen was readily apparent within and around hair follicles. Viral antigen was also evident in structures consistent with the sebaceous glands of hair follicles. As infectious virus can be detected on the surface of skin from EBOV infected postmortem NHPs, these data and the lack of robust epidermal infection suggests EBOV may utilize hair follicles as a mechanism for viral trafficking to the epidermal surface (cite NHP paper and hair follicle structure).

EBOV stimulates proinflammatory gene transcripts in several key tissues during infection (20, 28, 41, 42). Consistent with previous studies, we found liver exhibited upregulated proinflammatory gene transcripts without a corresponding increase in anti-inflammatory genes. Adipocytes also showed strong inflammatory responses (20). Interestingly, in ma-EBOV-infected visceral fat, both pro- and anti-inflammatory gene transcripts were upregulated at 5 days post-infection, suggesting a mixed inflammatory environment. Similarly, belly skin had a mixed inflammatory environment; although, the belly skin tissue may be influenced by the highly proinflammatory subcutaneous fat. The unique inflammatory profile of EBOV in skin remains unclear, as no significant changes in pro- or anti-inflammatory gene transcripts were detected in either thigh or back distal skin tissue.

Using rVSV/EBOV GP, we directly tested whether virus could infect skin and disseminate systemically in *Ifnar ^−/−^* mice. Removal of the stratum corneum by abrasion led to higher lethality, as evidenced by survival and infectious titers in key organs. Interestingly, despite skin providing mice with some protection from high viral inoculums, viral dissemination still occurred as determined by high EBOV-GP specific IgG antibody titers. This suggests skin could serve as a potential route for EBOV vaccination (43).

Axl is a key PS receptor that mediates EBOV infection of primary and immortalized human dermal fibroblasts and keratinocytes (5), while Tim-1 and Tim-4 also facilitate EBOV GP-mediated entry in various cell types (11, 36, 39). We expected mice lacking Axl expression to have restricted skin infection but instead found that *Tim-1 ^−/−^* mice exhibited lower viral titers in both epidermis and dermis. We also found *Tim-1 ^−/−^* mice had reduced viral dissemination from skin. This aligns with prior findings that Tim-1 contributes to EBOV pathogenesis by decreasing titers in the liver at later times during infection and promoting proinflammatory cytokine dysregulation (7, 31). Together, our results highlight the importance of skin as a key site of EBOV infection and dissemination, reinforcing its role in systemic EBOV pathogenesis.

---

**Supplemental Figure 1.**
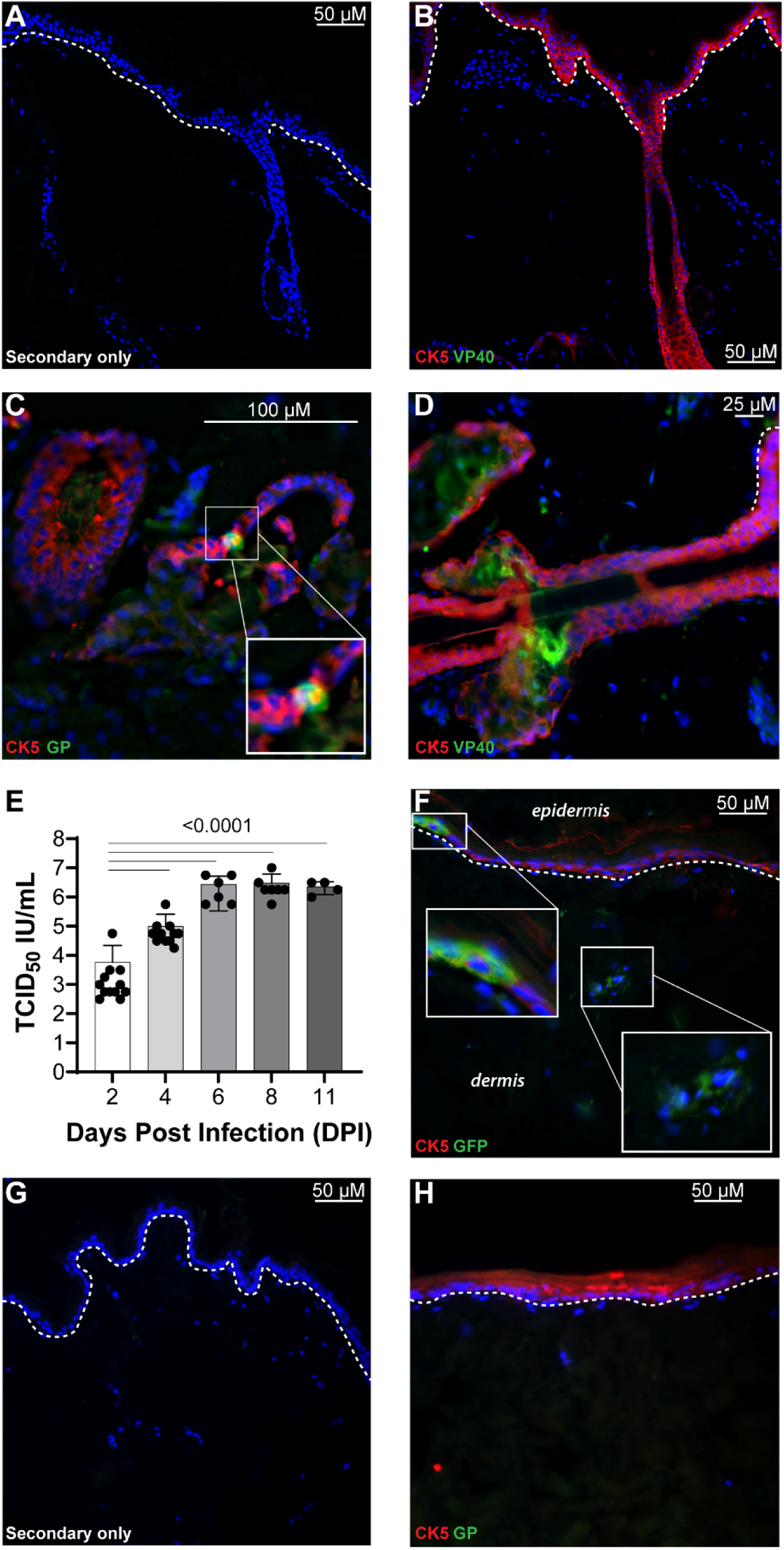
EBOV replicates and traffics to NHP skin at late times of systemic infection: NHP explant rVSV/EBOV GP infection and staining controls of NHP *in vivo* and explant tissues. A-B) Staining control examples used in *in vivo* NHP experiments described in Figure 1 for NHP skin tissue of infected skin tissue with secondary antibodies alone (**A**) and primary and secondary antibodies in mock-infected NHP skin (**B**). Additional FFPE sections showing hair follicle infection as described in Figure 1 were taken at (**C**) or contralateral to (**D**) the site of injection at either 6 (**C**) or 7 (**D**) days post infection. Sections were mounted in DAPI (blue) and stained with virus (EBOV GP or VP40 in green) and cell marker, CK5. **E-H)** 1 cm^2^ NHP skin explants from n=2 animals were generated from uninfected NHP skin. NHP skin explants were placed dermal side down on semipermeable transwell membranes and maintained at the air-liquid interface. Explants were infected via the basal media with 1.5×10^7^ IU of rVSV/EBOV GP in the presence of STAT1 inhibitor, B18R. Following overnight infection, explants were washed with DPBS twice to remove input and media was changed every other day up to 11 days post infection. **E)** Production of infectious virus in the explants was measured in the basal supernatants over time on Vero cells. Shown are 2 independent experiments with data expressed on a log_10_ scale as means ±SD. Ordinary one-way ANOVA, ****p<0.0001. **F-H)** NHP skin explants were fixed on day 8 post infection in 4% PFA and stained for viral antigen (GFP or GP, green) and keratinocyte marker (CK5, red) mounted in DAPI (blue). The dotted line indicates the epidermal-dermal junction. **F)** Viral antigen was detected in the dermis and colocalized with CK5+ keratinocytes as highlighted by insets. **G-H)** Staining control examples for NHP explant skin tissue of infected skin tissue with secondary antibodies alone (**G**) and primary and secondary antibodies in mock-infected skin explants (**H**). Entire images were enhanced by increasing brightness and decreasing contrast for visualization.

**Supplemental Figure 2.**
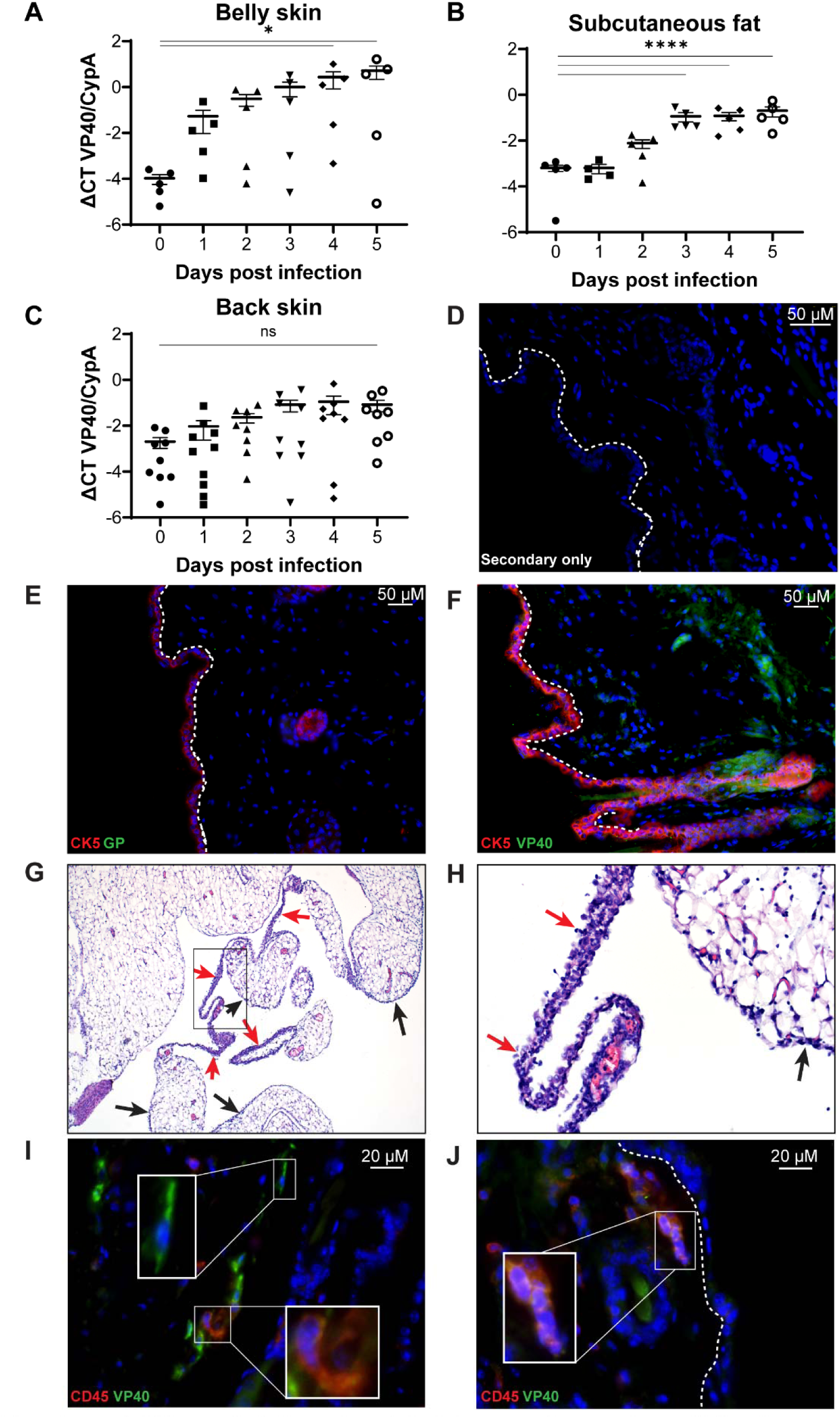
Adjacent and distal mouse tissues have distinct ma-EBOV kinetics and viral cell tropism: staining controls and additional images. C57BL6/N mice were infected and tissues were collected and processed as described in Figure 2. **A-C**) Viral load was assessed by RT-qPCR in the belly skin (**A**), subcutaneous fat (**B**), and back skin (**C**) over time. Viral load (VP40) was normalized to the house keeping gene cyclophilin-A (CypA). Data is expressed on a log_10_ scale as means ±SEM. Ordinary one-way ANOVA, *p<0.05 ****p<0.0001. **D, E**) Representative staining control images of infected day 5 thigh tissue with secondary antibody alone (**D**) and mock infected thigh tissue with both primary and secondary antibody staining (**E**). **F**) Additional FFPE sections stained with CK5 and VP40 show hair follicle infection as described in Figure 2 were taken from thigh at 5 days post infection. **G,H**) Hematoxylin and eosin (H&E) staining of visceral fat from day 5 infected mice. Increased cellularity is evident in visceral ligaments (red arrows) and patches of the lining visceral peritoneum (black arrows) (**G**). Inset from (**G**) demonstrates adipose targeted inflammation (arrows) of mononuclear cells and scattered neutrophils with evidence of necrosis (**H**). **I,J**) FFPE sections were taken from 4 days post infection in the thigh skin and stained for viral antigen (VP40, green) and CD45 (red). CD45-negative viral antigen-positive cells were present in the dermis and morphologically appear like fibroblasts (**I, top inset**). Small foci of CD45-positive and viral antigen-positive cells were evident (**I, bottom inset, J, inset**). Entire images were enhanced by increasing brightness and decreasing contrast for visualization.

**Supplemental Figure 3.**
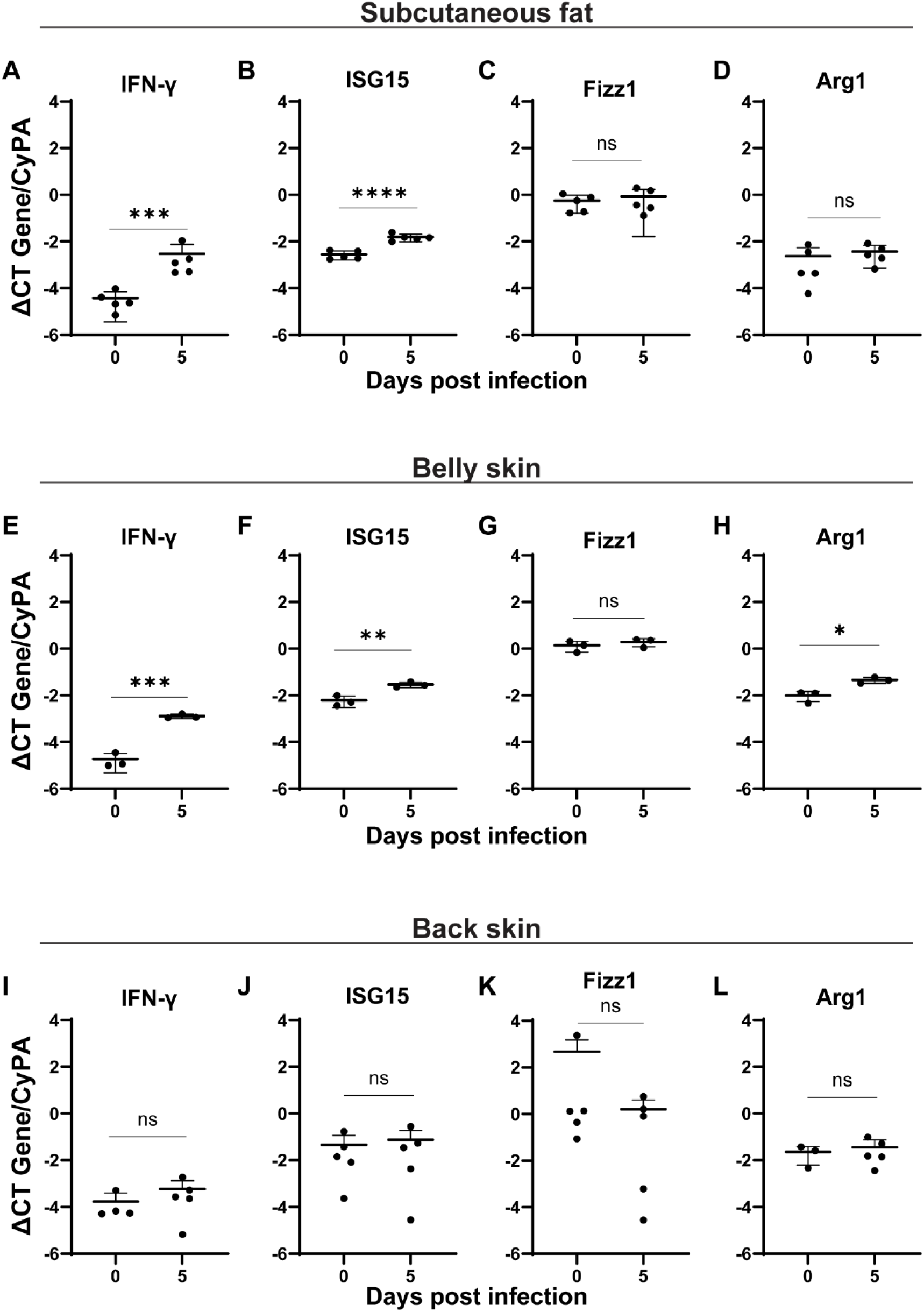
Inflammatory profiles of multiple mouse tissues infected with ma-EBOV: additional tissues. C57BL6/N mice were infected and tissues were collected and processed as described in Figure 2. Proinflammatory (IFN-γ and ISG15) and anti-inflammatory (Fizz1 and Arg1) gene transcripts were measured in the subcutaneous fat (**A-D**), belly skin (**E-H**), and back skin (**I-L**). Values were measured in samples taken at 0 and 5 days post infection and normalized to the house keeping gene cyclophilin-A (CypA). Data is expressed on a log_10_ scale as means ±SD. Student’s t test, **p<0.01 ***p<0.001 ****p<0.0001.

**Supplemental Figure 4.**
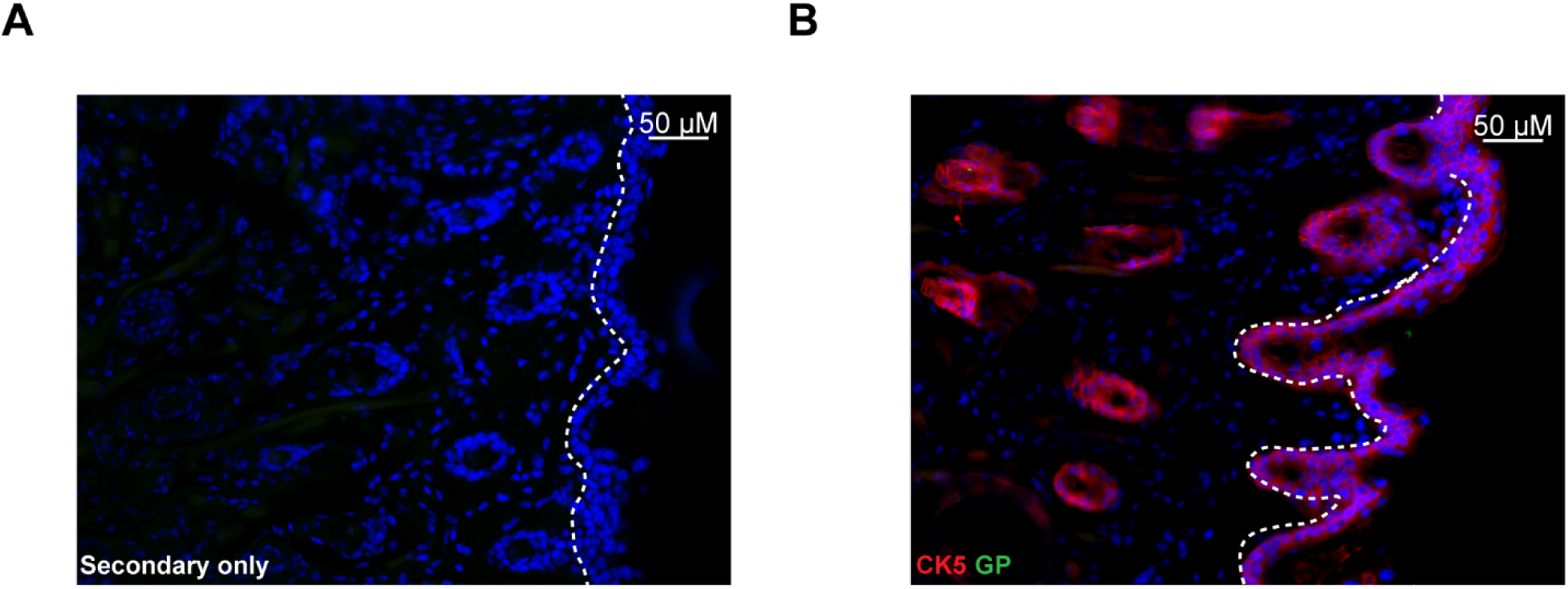
rVSV/EBOV GP mouse infections are similar to ma-EBOV: staining controls. Representative staining control images of infected day 3 cheek tissue with secondary antibody alone (**A**) and mock infected thigh tissue with both primary and secondary antibody staining (**B**).

**Supplemental Figure 5.**
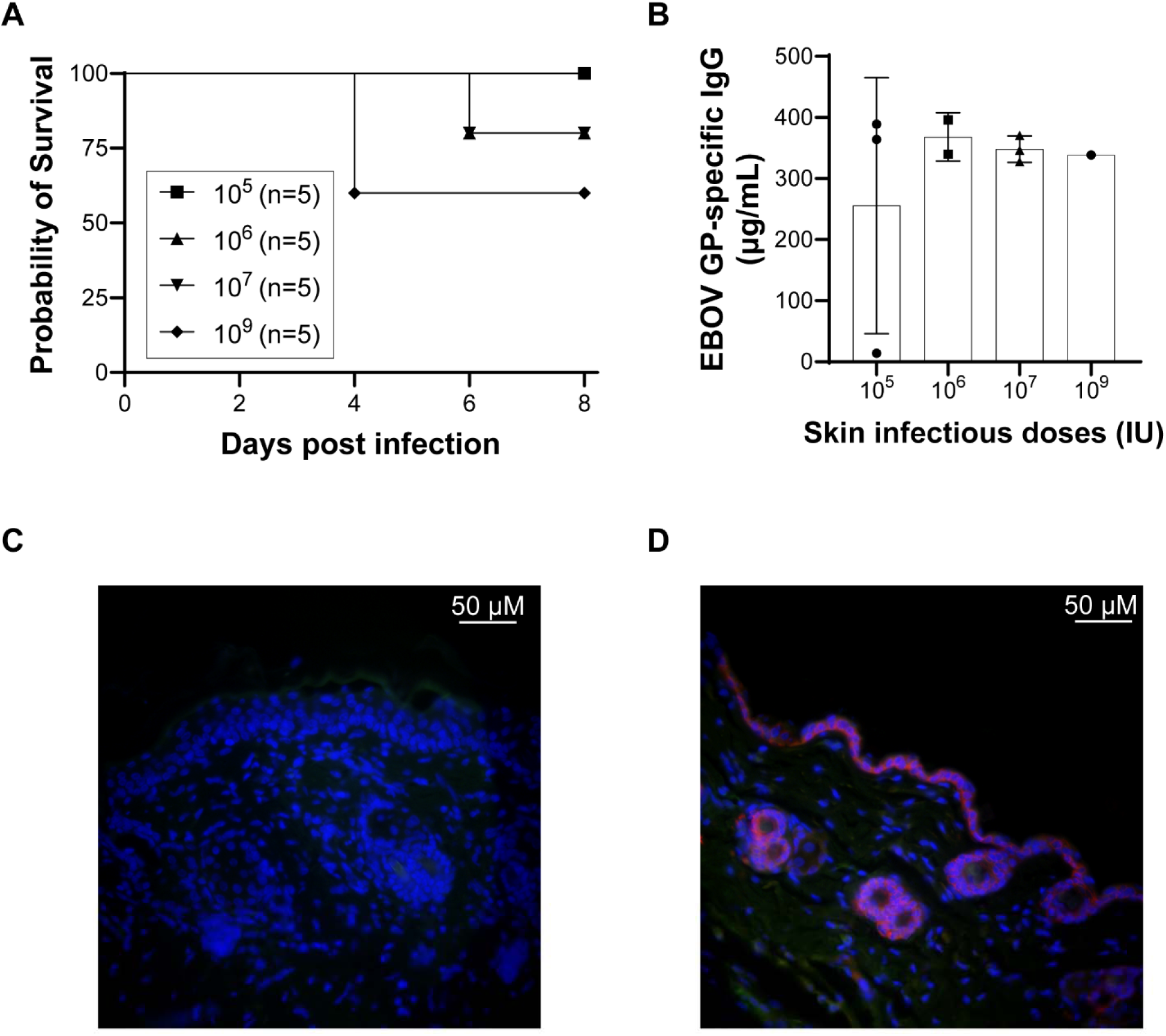
Direct infection of the skin epidermal surface in vivo results in viral dissemination and morbidity: Dose response survival curve, EBOV-GP antibody detection, and staining controls. 6-10 week old male *Ifnar ^−/−^* C57BL6/J mice were shaved on the back right flank and allowed to rest for five days. 10 μL of 10^5^ - 10^9^ IU of rVSV/EBOV GP was placed on the shaved skin after gentle abrasion (abraded). Survival was monitored and Log-rank test was performed (n=5 per group) (**A**) and anti-EBOV GP IgG antibody serum titers were measured in mice that survived to 21 days post infection (**B**). Representative staining control images of 24-hour post infection back tissue with secondary antibody alone (**C**) and mock infected thigh tissue with both primary and secondary antibody staining (**D**).

**Supplemental Figure 6.**
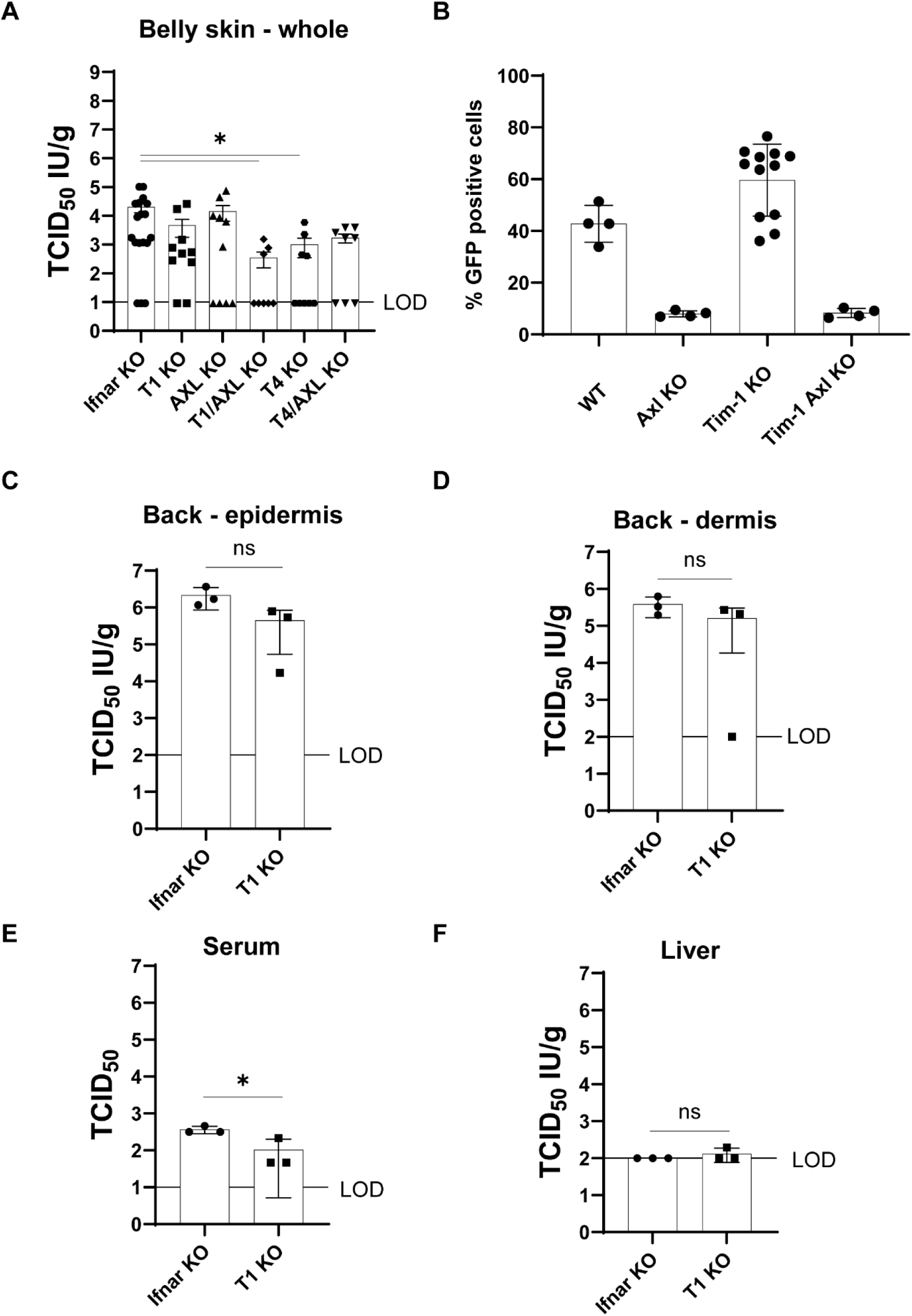
PS receptors mediate viral trafficking to and from skin: primary mouse fibroblast infection. **A**) 6-10 week old *Tim-1 ^−/−^ Ifnar ^−/−^* (n=11), *Axl ^−/−^ Ifnar ^−/−^* (n=10), *Tim-4 ^−/−^ Ifnar ^−/−^*(n=11), *Tim-1^−/−^* | *Axl ^−/−^ Ifnar ^−/−^* (n=10), *Tim-4^−/−^* | *Axl ^−/−^ Ifnar ^−/−^* (n=10), and *Ifnar ^−/−^* (n=17) C57BL6/J mice were shaved on the back right flank and allowed to rest for five days. All mice were infected with 500 IU of rVSV/EBOV GP and euthanized at 3 days post infection. Intact belly skin tissue was collected, homogenized, and titered on Vero E6 cells. Ordinary one-way ANOVA, *p<0.05. **B**) Back skin from uninfected *ifnar ^−/−^*, *Tim-1 ^−/−^ Ifnar ^−/−^, Axl ^−/−^ Ifnar ^−/−^,* and *Tim-1^−/−^* | *Axl ^−/−^ Ifnar ^−/−^* C57BL6/J mice. The dermis was embedded into a tissue culture treated plate with a scalpel. Primary fibroblasts outgrew from the embedded tissue and were passaged one time. Cells were plated in a 48-well plate at 100,000 cells per well and infected at an MOI of 10 with rVSV/EBOV GP. 48 hours post infection, cells were lifted with 0.25% trypsin and analyzed by flow cytometry for GFP. **C-F**) 6-10 week old *Tim-1 ^−/−^ Ifnar ^−/−^* (n=3) and *Ifnar ^−/−^* (n=3) C57BL6/J mice were shaved on the back right flank and allowed to rest for five days. 10 μL of 10^9^ IU of rVSV/EBOV GP was placed on the shaved skin following gentle abrasion. 24 hours after infection mice were euthanized and viral titers were measured in enzymatically processed back skin tissue to dissociate and separate the epidermis (**C**) and dermis (**D**). The epidermis and dermis were homogenized and titered on Vero E6 cells. Serum (**E**) and liver (**F**) were also collected or homogenized and tittered on Vero E6 cells. Statistics was determined by student’s T test on transformed values *p<0.05.

## REFERENCES

1. CDC. 2024. Outbreak History. Ebola. https://www.cdc.gov/ebola/outbreaks/index.html. Retrieved 10 March 2025.

2. Ten health issues WHO will tackle this year. https://www.who.int/news-room/spotlight/ten-threats-to-global-health-in-2019. Retrieved 4 March 2025.

3. Ansari AA. 2014. Clinical features and pathobiology of Ebolavirus infection. Journal of Autoimmunity 55:1–9.

4. Ericson AD, Claude KM, Vicky KM, Lukaba T, Richard KO, Hawkes MT. 2020. Detection of Ebola virus from skin ulcers after clearance of viremia. J Clin Virol 131:104595.

5. Prescott J, Bushmaker T, Fischer R, Miazgowicz K, Judson S, Munster VJ. 2015. Postmortem Stability of Ebola Virus. Emerg Infect Dis 21:856–859.

6. Messingham KN, Richards PT, Fleck A, Patel RA, Djurkovic M, Elliff J, Connell S, Crowe TP, Munoz Gonzalez J, Gourronc F, Dillard JA, Davey RA, Klingelhutz A, Shtanko O, Maury W. 2025. Multiple cell types support productive infection and dynamic translocation of infectious Ebola virus to the surface of human skin. Science Advances 11:eadr6140.

7. Bohan D, Maury W. 2021. Enveloped RNA virus utilization of phosphatidylserine receptors: Advantages of exploiting a conserved, widely available mechanism of entry. PLOS Pathogens 17:e1009899.

8. Automation of Infectious Focus Assay for Determination of Filovirus Titers and Direct Comparison to Plaque and TCID50 Assays. https://www.mdpi.com/2076-2607/9/1/156. Retrieved 3 April 2025.

9. Hierholzer JC, Killington RA. 1996. 2 -Virus isolation and quantitation, p. 25–46. In Mahy, BW, Kangro, HO (eds.), Virology Methods Manual. Academic Press, London.

10. Alfson KJ, Goez-Gazi Y, Gazi M, Staples H, Mattix M, Ticer A, Klaffke B, Stanfield K, Escareno P, Keiser P, Griffiths A, Chou Y-L, Niemuth N, Meister GT, Cirimotich CM, Carrion R. 2021. Development of a Well-Characterized Rhesus Macaque Model of Ebola Virus Disease for Support of Product Development. 3. Microorganisms 9:489.

11. Brunton B, Rogers K, Phillips EK, Brouillette RB, Bouls R, Butler NS, Maury W. 2019. TIM-1 serves as a receptor for ebola virus in vivo, enhancing viremia and pathogenesis. PLoS Neglected Tropical Diseases 13:e0006983.

12. Rogers KJ, Shtanko O, Vijay R, Mallinger LN, Joyner CJ, Galinski MR, Butler NS, Maury W. 2020. Acute Plasmodium Infection Promotes Interferon-Gamma-Dependent Resistance to Ebola Virus Infection. Cell Reports 30:4041–4051.e4.

13. Sakurai Y, Kolokoltsov AA, Chen C-C, Tidwell MW, Bauta WE, Klugbauer N, Grimm C, Wahl-Schott C, Biel M, Davey RA. 2015. Ebola virus. Two-pore channels control Ebola virus host cell entry and are drug targets for disease treatment. Science 347:995–998.

14. Siragam V, Wong G, Qiu X-G, Siragam V, Wong G, Qiu X-G. 2018. Animal models for filovirus infections. Zoological Research, 2018, Vol 39, Issue 1, Pages: 15–24 39:15–24.

15. Zaki SR, Goldsmith CS. 1999. Pathologic features of filovirus infections in humans. Curr Top Microbiol Immunol 235:97–116.

16. Gire SK, Goba A, Andersen KG, Sealfon RSG, Park DJ, Kanneh L, Jalloh S, Momoh M, Fullah M, Dudas G, Wohl S, Moses LM, Yozwiak NL, Winnicki S, Matranga CB, Malboeuf CM, Qu J, Gladden AD, Schaffner SF, Yang X, Jiang P-P, Nekoui M, Colubri A, Coomber MR, Fonnie M, Moigboi A, Gbakie M, Kamara FK, Tucker V, Konuwa E, Saffa S, Sellu J, Jalloh AA, Kovoma A, Koninga J, Mustapha I, Kargbo K, Foday M, Yillah M, Kanneh F, Robert W, Massally JLB, Chapman SB, Bochicchio J, Murphy C, Nusbaum C, Young S, Birren BW, Grant DS, Scheiffelin JS, Lander ES, Happi C, Gevao SM, Gnirke A, Rambaut A, Garry RF, Khan SH, Sabeti PC. 2014. Genomic surveillance elucidates Ebola virus origin and transmission during the 2014 outbreak. Science 345:1369–1372.

17. Millar SE. 2002. Molecular Mechanisms Regulating Hair Follicle Development. Journal of Investigative Dermatology 118:216–225.

18. Bray M, Davis K, Geisbert T, Schmaljohn C, Huggins J. 1999. A Mouse Model for Evaluation of Prophylaxis and Therapy of Ebola Hemorrhagic Fever. The Journal of Infectious Diseases 179:S248–S258.

19. Gibb TR, Bray M, Geisbert TW, Steele KE, Kell WM, Davis KJ, Jaax NK. 2001. Pathogenesis of Experimental Ebola Zaire Virus Infection in BALB/c Mice. Journal of Comparative Pathology 125:233–242.

20. Gourronc FA, Rebagliati MR, Kramer-Riesberg B, Fleck AM, Patten JJ, Geohegan-Barek K, Messingham KN, Davey RA, Maury W, Klingelhutz AJ. 2022. Adipocytes are susceptible to Ebola Virus infection. Virology 573:12–22.

21. Martines RB, Ng DL, Greer PW, Rollin PE, Zaki SR. 2015. Tissue and cellular tropism, pathology and pathogenesis of Ebola and Marburg viruses. The Journal of Pathology 235:153–174.

22. Liu X, Speranza E, Muñoz-Fontela C, Haldenby S, Rickett NY, Garcia-Dorival I, Fang Y, Hall Y, Zekeng E-G, Lüdtke A, Xia D, Kerber R, Krumkamp R, Duraffour S, Sissoko D, Kenny J, Rockliffe N, Williamson ED, Laws TR, N’Faly M, Matthews DA, Günther S, Cossins AR, Sprecher A, Connor JH, Carroll MW, Hiscox JA. 2017. Transcriptomic signatures differentiate survival from fatal outcomes in humans infected with Ebola virus. Genome Biol 18:4.

23. McElroy AK, Mühlberger E, Muñoz-Fontela C. 2018. Immune barriers of Ebola virus infection. Current Opinion in Virology 28:152–160.

24. Messaoudi I, Amarasinghe GK, Basler CF. 2015. Filovirus pathogenesis and immune evasion: insights from Ebola virus and Marburg virus. Nat Rev Microbiol 13:663–676.

25. Eisfeld AJ, Halfmann PJ, Wendler JP, Kyle JE, Burnum-Johnson KE, Peralta Z, Maemura T, Walters KB, Watanabe T, Fukuyama S, Yamashita M, Jacobs JM, Kim Y-M, Casey CP, Stratton KG, Webb-Robertson B-JM, Gritsenko MA, Monroe ME, Weitz KK, Shukla AK, Tian M, Neumann G, Reed JL, van Bakel H, Metz TO, Smith RD, Waters KM, N’jai A, Sahr F, Kawaoka Y. 2017. Multi-platform ‘Omics Analysis of Human Ebola Virus Disease Pathogenesis. Cell Host Microbe 22:817–829.e8.

26. Escudero-Pérez B, Volchkova VA, Dolnik O, Lawrence P, Volchkov VE. 2014. Shed GP of Ebola Virus Triggers Immune Activation and Increased Vascular Permeability. PLOS Pathogens 10:e1004509.

27. Martins K, Cooper C, Warren T, Wells J, Bell T, Raymond J, Stuthman K, Benko J, Garza N, van Tongeren S, Donnelly G, Retterer C, Dong L, Bavari S. 2015. Characterization of Clinical and Immunological Parameters During Ebola Virus Infection of Rhesus Macaques. Viral Immunology 28:32–41.

28. Hensley LE, Young HA, Jahrling PB, Geisbert TW. 2002. Proinflammatory response during Ebola virus infection of primate models: possible involvement of the tumor necrosis factor receptor superfamily. Immunology Letters 80:169–179.

29. Bosio CM, Aman MJ, Grogan C, Hogan R, Ruthel G, Negley D, Mohamadzadeh M, Bavari S, Schmaljohn A. 2003. Ebola and Marburg Viruses Replicate in Monocyte-Derived Dendritic Cells without Inducing the Production of Cytokines and Full Maturation. The Journal of Infectious Diseases 188:1630–1638.

30. Sanchez A, Lukwiya M, Bausch D, Mahanty S, Sanchez AJ, Wagoner KD, Rollin PE. 2004. Analysis of Human Peripheral Blood Samples from Fatal and Nonfatal Cases of Ebola (Sudan) Hemorrhagic Fever: Cellular Responses, Virus Load, and Nitric Oxide Levels. Journal of Virology 78:10370–10377.

31. Richards PT, Briseño JAA, Brunton BA, Maury W. 2025. In Vivo Investigation of Filovirus Glycoprotein-Mediated Infection in a BSL2 Setting. Methods Mol Biol 2877:183–198.

32. Rhein BA, Powers LS, Rogers K, Anantpadma M, Singh BK, Sakurai Y, Bair T, Miller-Hunt C, Sinn P, Davey RA, Monick MM, Maury W. 2015. Interferon-γ Inhibits Ebola Virus Infection. PLoS Pathogens 11.

33. Victory KR, Coronado F, Ifono SO, Soropogui T, Dahl BA, Centers for Disease Control and Prevention (CDC). 2015. Ebola transmission linked to a single traditional funeral ceremony - Kissidougou, Guinea, December, 2014-January 2015. MMWR Morb Mortal Wkly Rep 64:386–388.

34. Alvarez CP, Lasala F, Carrillo J, Muñiz O, Corbí AL, Delgado R. 2002. C-Type Lectins DC-SIGN and L-SIGN Mediate Cellular Entry by Ebola Virus in cis and in trans. Journal of Virology 76:6841–6844.

35. Shimojima M, Takada A, Ebihara H, Neumann G, Fujioka K, Irimura T, Jones S, Feldmann H, Kawaoka Y. 2006. Tyro3 Family-Mediated Cell Entry of Ebola and Marburg Viruses. Journal of Virology 80:10109–10116.

36. Kondratowicz AS, Lennemann NJ, Sinn PL, Davey RA, Hunt CL, Moller-Tank S, Meyerholz DK, Rennert P, Mullins RF, Brindley M, Sandersfeld LM, Quinn K, Weller M, McCray PB, Chiorini J, Maury W. 2011. T-cell immunoglobulin and mucin domain 1 (TIM-1) is a receptor for Zaire Ebolavirus and Lake Victoria Marburgvirus. Proceedings of the National Academy of Sciences 108:8426–8431.

37. Shimojima M, Ikeda Y, Kawaoka Y. 2007. The Mechanism of Axl-Mediated Ebola Virus Infection. The Journal of Infectious Diseases 196:S259–S263.

38. Richard AS, Zhang A, Park S-J, Farzan M, Zong M, Choe H. 2015. Virion-associated phosphatidylethanolamine promotes TIM1-mediated infection by Ebola, dengue, and West Nile viruses. Proceedings of the National Academy of Sciences 112:14682–14687.

39. Rhein BA, Brouillette RB, Schaack GA, Chiorini JA, Maury W. 2016. Characterization of Human and Murine T-Cell Immunoglobulin Mucin Domain 4 (TIM-4) IgV Domain Residues Critical for Ebola Virus Entry. Journal of Virology 90:6097–6111.

40. Spengler JR, Welch SR, Ritter JM, Harmon JR, Coleman-McCray JD, Genzer SC, Seixas JN, Scholte FEM, Davies KA, Bradfute SB, Montgomery JM, Spiropoulou CF. 2023. Mouse models of Ebola virus tolerance and lethality: characterization of CD-1 mice infected with wild-type, guinea pig-adapted, or mouse-adapted virus. Antiviral Research 210:105496.

41. Basler CF. 2015. Innate immune evasion by filoviruses. Virology 10.1016/j.virol.2015.03.030.

42. Basler CF, Amarasinghe GK. 2009. Evasion of interferon responses by ebola and marburg viruses. Mary Ann Liebert, Inc. 10.1089/jir.2009.0076.

43. Romanyuk A, Wang R, Marin A, Janus BM, Felner EI, Xia D, Goez-Gazi Y, Alfson KJ, Yunus AS, Toth EA, Ofek G, Carrion R, Prausnitz MR, Fuerst TR, Andrianov AK. 2022. Skin Vaccination with Ebola Virus Glycoprotein Using a Polyphosphazene-Based Microneedle Patch Protects Mice against Lethal Challenge. J Funct Biomater 14:16.

